# Targeting activated macrophages intracellular milieu to augment anti-inflammatory drug potency

**DOI:** 10.1101/2021.06.22.449368

**Authors:** Virgínia M. Gouveia, Loris Rizzello, Bruno Vidal, Claudia Nunes, Alessandro Poma, Ciro Lopez-Vasquez, Edoardo Scarpa, Sebastian Brandner, António Oliveira, João E. Fonseca, Salette Reis, Giuseppe Battaglia

**Affiliations:** Department of Chemistry, University College London, WC1H 0AJ London, UK; Institute of Physics of Living Systems, University College London WC1H 0AJ London, UK; SomaServe Ltd, Babraham Research Campus, CB22 3AT Cambridge, UK; LAQV, REQUIMTE, Department of Chemical Sciences, Faculty of Pharmacy, University of Porto, 4050-313 Porto, Portugal; Abel Salazar Biomedical Sciences Institute, University of Porto, 4050-313 Porto, Portugal; Institute for Bioengineering of Catalonia (IBEC), The Barcelona Institute of Science and Technology, 08028 Barcelona, Spain; Department of Pharmaceutical Sciences, University of Milan, 20133 Milan, Italy; National Institute of Molecular Genetics (INGM), 20122 Milan, Italy; Division of Biomaterials and Tissue Engineering, Eastman Dental Institute, Royal Free Hospital, UCL Medical School, NW3 2PF London, UK; Department of Neurodegenerative Disease, Queen Square Institute of Neurology, University College London, WC1N 3BG London, UK; Rheumatology Research Unit, Institute of Molecular Medicine – IMM João Lobo Antunes, Faculty of Medicine, University of Lisbon, 1649-028 Lisbon, Portugal; Serviço de Reumatologia, Centro Hospitalar Universitário Lisboa Norte, Centro Académico de Medicina de Lisboa, 1649-028 Lisbon, Portugal; Catalan Institution for Research and Advanced Studies (ICREA), 08010 Barcelona, Spain

## Abstract

We present pH-responsive phosphorylcholine polymersomes ability to target activated macrophages via scavenger receptors, enter them via endocytosis, and escape from early endosomes enabling the intra-cellular drug delivery. Using an arthritis experimental model and the gold standard disease modifying anti-rheumatic drug, methotrexate, we prove that polymersomes augments therapeutic efficacy, while minimizing the off-target effect. First, we demonstrate the selective accumulation of polymersomes within the inflamed synovial tissues and cells, including macrophages. Second, we show the beneficial therapeutic effect of methotrexate loaded polymersomes in preventing both joint inflammation and further damage. Hence, we prove the therapeutic potential of polymersomes in enhancing the complete prevention of arthritis progression, which makes it a promising nanotherapy for arthritis treatment as well as other inflammatory disorders.

**Teaser:** We show that the effective targeting and delivery of drugs to the main inflammation actors, the macrophages, enhances arthritis therapy.

## Introduction

Macrophages are one of the most important immune cells and as such, they are present in most organs adapting to the local environment patrolling it from external attack. Macrophages are the main modulators of inflammation and the primary immune response to cellular damage. They have both pro- or anti-inflammatory actions depending on their surrounding feedback, such process is referred to as polarization and often macrophages differentiate plastically from a M1 stage (pro) at the initial stages to M2 (anti) during the resolution stages of inflammation (*1*). Macrophage polarization is a critical step in both resolving inflammation and promoting, tissue repair, and any variations in its sequences give rise to immunological disorders including, autoimmune and metabolic diseases, chronic inflammation, and even cancer (*1*). One of the most common of such disorders, is Rheumatoid arthritis (RA), a chronic inflammatory immune-mediated disease that affects nearly 1% of the population worldwide (*2–4*). This arthritic disorder is characterized by the progressive inflammation of the synovial tissue that causes loss of joint functionality (*3–6*). Severe disease progression can lead to cartilage and bone damage and irreversible destruction of joints (*4, 6*). RA pathogenesis involves the interplay of both innate and adaptive immune responses (*6–8*). The influx of immune cells to the synovium and subsequent interacting cascades of pro-inflammatory cytokines causes synovitis (*6–8*). Synovium inflammation increases vascular permeability, which in turn leads to synovial hyperplasia and formation of inflamed pannus tissue (*9, 10*). More importantly, local synovial cellular interactions initiate and maintain the inflammatory process through the activation of macrophages and later on synovial fibroblasts (also known as synoviocytes) (*6–8*). Both cell types are found in abundance in the synovium and play a central role in RA pathogenesis by actively driving the perpetuation of immune and inflammatory responses and further damage of joint tissue (*6, 7, 11, 12*). Macrophages induce the secretion of several pro-inflammatory cytokines, such as tumor necrosis factor- (TNF), interleukin (IL)1β and IL6, and chemokines, such as CXCL8 (*i.e.* IL8) and CCL3 (also known as macrophage inflammatory protein 1α) (*6–8, 12*). Through cascades of cytokine-mediated signalling pathways, synovial macrophages activate other cells found within the synovium that are involved in the disease progression (*6–8, 11, 12*). Among them, synoviocytes inflammatory phenotype activation stimulates the production of more pro-inflammatory cytokines and chemokines, thus perpetuating synovial inflammation. These cells are important in the maintenance of the normal stromal synovial environment of joints through the secretion of a range of extracellular matrix components (*6, 11, 13–15*). Activated synoviocytes increase the secretion of tissue-degrading enzymes, such as the matrix metalloproteinases (MMPs) that are involved in the degradation of cartilage (*6, 11, 13–15*). In addition, the synoviocytes secrete the receptor activator of nuclear factor-κB (NF-κB) ligand (RANKL), which, in turn, induce the differentiation and proliferation of osteoclasts that are mainly responsible for bone erosion (*6, 11, 13–15*). Moreover, synoviocytes play an important role in regulating the disease progression through the secretion of vascular endothelial growth factors associated with angiogenesis (*9, 10, 15–18*). The enhanced synovial vascularization and prominent angiogenesis endures synovitis and facilitates the access of synoviocytes to the bloodstream, hence increasing the dissemination of arthritis to other unaffected joints (*9, 15–17, 19*).

RA is a serious chronic unremitting health condition, and despite the advances on treatment modalities, which focus mostly on the inflammation, it has, (*20–22*). The management of RA relies on the early treatment with disease-modifying anti-rheumatic drugs (DMARDs) to relief the symptoms and cease the progression of synovial inflammation (*3, 4, 20, 22*). Thereby, early treatment intervention is crucial to slow down the irreversible joint damage (*23*). Methotrexate (MTX) is the gold standard DMARD in RA and is used also in other immune mediated chronic inflammatory diseases currently (*3, 20, 22*). Different mechanisms of action are involved in the therapeutic effectiveness of this drug (*24–26*). Namely, MTX adenosine-mediated anti-inflammatory and immunosuppressive effects regulate the production of cellular adhesion and pro-inflammatory molecules involved in synovial and systemic inflammation (*24–26*). Despite the therapeutic efficacy of MTX in clinic, a significant number of patients discontinue the therapy, mostly due to adverse effects and intolerance (*3, 22, 24, 27–30*).

In the last 15 years, several nanomedicines have been developed to improve MTX selectivity and on-site delivery in inflamed synovial tissues to overcome the limitations related to the use of DMARD for RA treatment (*4, 31, 32*). In this study, we assessed the use of pH-responsive polymersomes made of poly(2-(methacryloyloxy) ethyl phosphorylcholine – poly(2-(diisopropylamino)ethyl methacrylate (PMPC–PDPA) as a drug delivery system to precisely target and treat synovial inflammation. The potential of this nanomedicine is founded on the polymersomes building blocks: PMPC endows cell targeting ability, via scavenger receptors (SR) class B type 1 and 3, and then cell entry along the endocytic pathway; whereas the PDPA pH responsiveness bestows the drugs release from early endosomes to the cell cytosol (*33–40*).

*In vitro* and *in vivo* findings support the potential of pH-responsive PMPC-PDPA polymersomes (subsequently designated psomes) to ensure the stability and selective delivery of MTX within inflamed joint synovium, macrophages and synoviocytes. We prove that MTX-loaded psomes efficiently shut down the immune-mediated inflammatory response *in vitro*. Additionally, we studied the biodistribution, biocompatibility and the therapeutic impact of psomes in the progression of chronic synovial inflammation in well-established adjuvant-induced arthritis animal model. Daily intraperitoneal administration of MTX-loaded psomes is highly effective in the inhibition of the inflammatory arthritic signs and in suppressing both local synovial and systemic inflammation. Remarkably, psomes alone can abrogate arthritic inflammation, as well as, the progression of systemic inflammation, in the same model of arthritis. In conclusion, we demonstrate the beneficial therapeutic effects of psomes to control the disease inflammatory activity and to hinder arthritis progression. Hence, establishing their potential to be used in the clinic for the treatment of RA.

## Results

### Psomes as suitable drug delivery system

Formulations of psomes (i.e. PMPC-PDPA polymersomes) and MTX-loaded psomes were successfully produced by pH-switch self-assembling methodology and then characterised in terms of size distribution and morphology. Dynamic light scattering (DLS) measurements revealed that both formulations presented unimodal size distribution profiles with hydrodynamic diameters (D_h_) of 93 ± 13 nm and 96 ± 11 nm, respectively. Transmission electron microscopy (TEM) characterisation also confirmed the spherical morphology of vesicles with a uniform size distribution of about 90 nm (**Fig. S1**). Likewise, TEM micrographs of both Cy5- and Cy7-psomes produced via solvent-switch present a homogeneous distribution of spherical polymersomes. The encapsulation of MTX was performed via modification of the pH-switch method. The drug was solubilised in a NaOH basic solution, rather than in the polymer solution at pH 2.0 before the self-assembly process. High- performance liquid chromatography (HPLC) quantifications confirmed the high drug loading capacity of psomes towards the MTX (up to 18 ± 2.1 wt%; **Fig. S2**). This confirms the efficacy of the solvent switch method to enable a higher drug loading, possibly due to the affinity of MTX for the polymer during the self-assembly process. The partition coefficient value at pH 7.4 (log D of 2.2 ± 0.1; **Fig. S3**) confirmed the affinity of MTX with the polymeric phase of the water/ polymersome system.

In vitro drug release studies were carried out overtime under physiologic and acidic pH conditions at 37°C. The burst drug release profile observed at pH 5.0, with the complete release of MTX within 1 hour incubation, confirmed psomes responsiveness to acidic pH in the intracellular endosomal compartments.), whereas we observed a more sustained drug release over time at pH 6.5 and negligible at pH 7.4, even after 100 hurs (**Fig. S4**). Drug release kinetic analyses based on the regression coefficient (r2) revealed that the best fitting model for pH 5.0 and pH 6.5 profiles was obtained by the Zero-Order and Higuchi models (**Table S1**). These models suggest that the MTX release occurs through a controlled diffusion mechanism (**Table S2**). In addition, and as expected, there was no best-fitted model for the release profile at pH 7.4 and practically negligible drug release was observed throughout 100 hours (**Fig. S4**). These results demonstrate the effectiveness of psomes as a nanomedicine that ensures the loading of MTX and its on-site release.

### Psomes biodistribution in AIA rats

The biodistribution of psomes in the blood stream for 24 hours was assessed to understand the pharmacokinetic behaviour and stability in vivo. Two experimental groups of Wistar rats (n=3), one healthy and other with adjuvant-induced arthritis (AIA), received an intraperitoneal (i.p.) injection of Cy7-psomes (10 mg/kg body weight). Blood samples were then collected by tail vein puncture at different time points. The blood plasma concentration profile ofinjected Cy7-psomes, in both healthy and arthritic animal models, had two phases – absorption and elimination –, characteristic of intraperitoneal (i.p.) administration (**Fig. 1A**, **Fig. S5A**). The absorption phase occurred within 2 hours, reaching a maximum peak of psome concentration in the plasma of 70 ± 23 μg/ml (**Fig. 1A**), followed by its elimination from the blood stream. We then analysed the pharmacokinetic parameters of the Cy7-psomes in the plasma after i.p. injection in healthy and AIA Wistar rats (**Fig. 1B**). The half-life (t_1/2_) of Cy7-psomes was calculated by one phase decay analysis of the elimination phase (dot lines; **Fig. 1A**). Despite the similarity of both plasma concentration-time profiles in the healthy and the AIA Wistar rats (**Fig. 1A**; **Fig. S5A**), pharmacokinetic analysis resulted in a longer t_1/2_ and 2.5 times slower elimination rate (K_e_) in AIA rats relatively to healthy ones (**Fig. 1B**). Also, the area under the curve (AUC) for the AIA profile was greater than for healthy group (**Fig. 1B**). These results suggest a longer circulation time of psomes in the bloodstream of AIA rats. Additionally, we studied the biodistribution of Cy7-psomes in various organs (spleen, kidneys, liver and heart) in both healthy and arthritic Wistar rats. After 24 hours from Cy7-psomes i.p. injection, all animals were euthanised and the organs were removed and wet weighed. The percentage of injected dose (%ID) per gram of tissue was then calculated by correlating the intensity of fluorescence of Cy7-psomes measured to the total weight of each organ. The analysis revealed high amounts of Cy7-psomes in the spleen of both groups (4.7 ± 1.2 and 3.0 ± 1.2 % for healthy and AIA, respectively; **Fig. S5B**). Conversely, the injected dose of Cy7-psomes that accumulated per gram of liver was only 1.0 ± 0.1 and 1.2 ± 0.1 % for healthy and AIA rats, respectively (**Fig. 1C**). Similarly, the amount of Cy7-psomes detected in heart and kidneys was negligible (**Fig. 1C**). Besides the distribution in the main organs, it was important to assess psomes distribution in the paws of both healthy and arthritic animals in order to confirm the nanocarrier ability to preferentially target the areas of synovial inflammation. IVIS imaging analysis revealed that no fluorescence intensity was detected in the paws of healthy rats oppositely to the AIA group (**Fig. 1D**). Interestingly, we also noticed that the fluorescence signal detected in the paws of arthritic rats was dependent on the degree of joint swelling/inflammation score (AIA section: increase from left to right; **Fig. 1D**). Normalisation of the fluorescence intensity signal to the vehicle treated control highlighted the significant differences between the healthy and AIA group of animals (**Fig. 1E**). Thereby, we confirm that psomes ably accumulate in the inflamed joints of AIA rats.

**Fig. 1.**
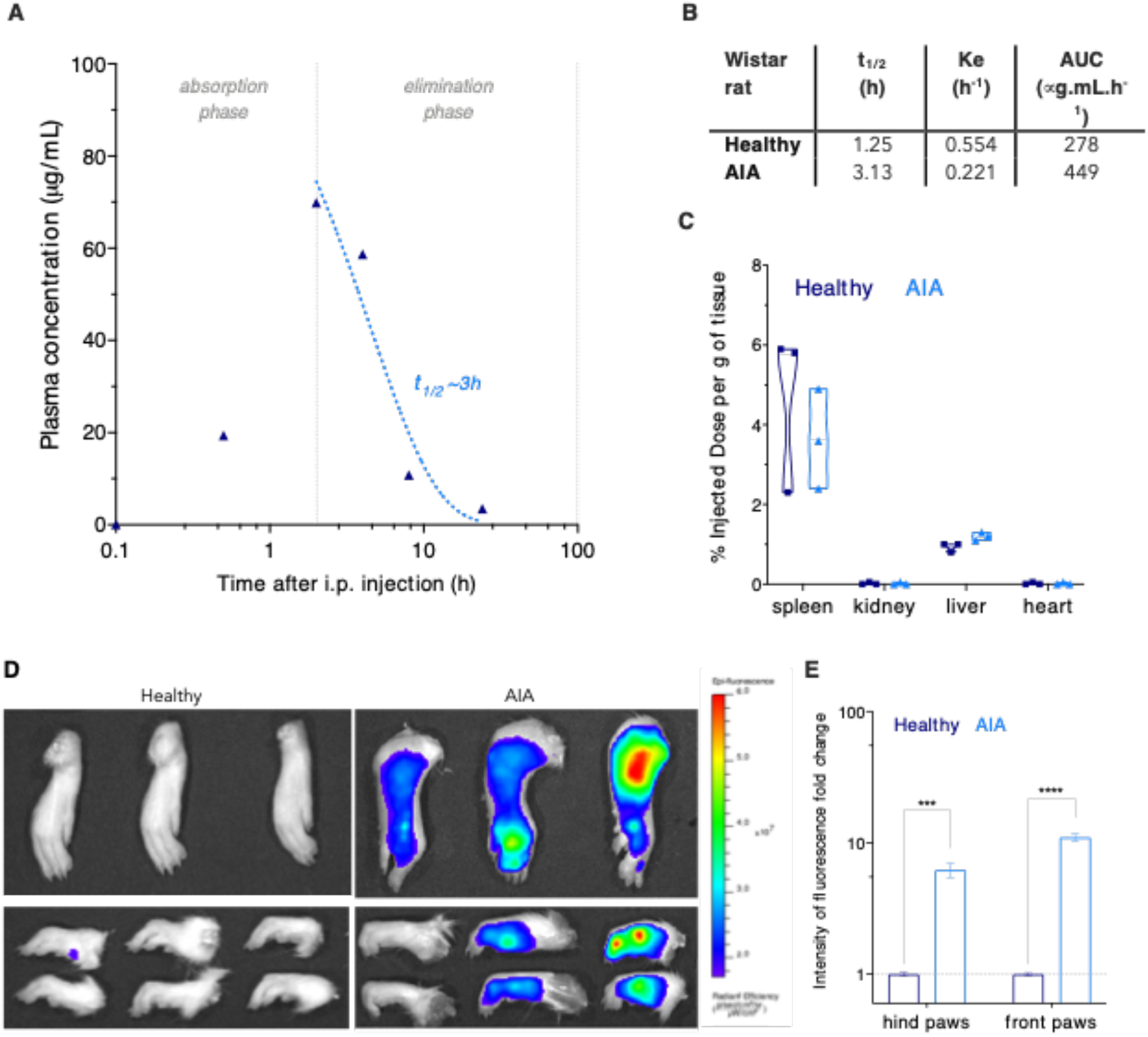
(**A**) Plasma concentration-time profile of Cy7-psomes after i.p. injection in AIA Wistar rats. (**B**) Pharmacokinetic parameters – terminal half-life (t_1/2_), elimination rate (K_e_), area under the curve (AUC) – regarding the plasma concentration-time profiles in healthy and AIA Wistar rats. (**C**) The injected dose (%ID) of Cy7-psomes per gram of organ tissue weight after 24 hours i.p. injection in healthy and AIA Wistar rats. (**D**) IVIS images of the right hind (top) and front (bottom) paws of healthy and AIA rats. (**E**) Fold change fluorescence normalised to the vehicle-treated control group. Data express the mean ± SEM (n=3 per experimental group). The differences relative to healthy group were statistically significant for *p<0.05, **p<0.01, ***p<0.001 and ****p<0.0001.

### Psomes enhanced uptake by macrophages and synoviocytes

The *in vitro* cellular uptake of psomes was assessed in non- or activated macrophages and synoviocytes using live confocal laser scanning microscopy (CLSM; **Fig. 2A-D**). CLSM imaging analysis of macrophages revealed that the normalised fluorescence intensity signal of Cy5-psomes constantly increased in a time-dependent manner up to 10-hours after incubation, followed by a plateau (**Fig. 2E**). Conversely, CLSM imaging on activated synoviocytes resulted in a triphasic cell uptake profile (**Fig. 2F**). First, there was an increase in the internalisation of Cy5-psomes, followed by a plateau of 5 hours and another increase of fluorescence intensity signal (of almost 100%). Oppositely, the uptake profile in non-activated synoviocytes suggest a slow internalisation of psomes overtime reaching a plateau after 1 hour of incubation (**Fig. 2F**). Both internalisation profiles suggest that there is a rapid and higher uptake of Cy5-psomes upon activation of both cell types comparing with the non-activated ones over the 48 hours (**Figure 2E-F**). Additional imaging analyses suggested that the fluorescence intensity of Cy5-psomes in LPS-activated macrophages was higher than in TNFα-activated synoviocytes (**Figure 2E-F**). This is not only due to the intrinsic professional phagocytic nature of macrophages (*41, 42*), but also possibly to the inherent binding affinity of the PMPC building block to the family of class B scavenger receptors, including type B1 and B3 (*33–36*). These surface cell receptors are expressed by macrophages and synoviocytes (*39, 40, 43, 44*). We then used western blot analysis to investigate the protein expression levels of these cell surface receptors followed by activation (**Fig. S7A**). Results revealed that SRB1 and SRB3 (commonly known as CD36) expression levels in activated macrophages increased by 2- and 4-fold, respectively, compared to non-activated cells (p<0.05 and p<0.01, respectively; **Fig. S7B**). Conversely, the activation of synoviocytes significantly boosted the expression of CD36 (p<0.0001 *versus* the non-activated control; **Fig. S7B**). However, the expression of SRB1 was only slightly increased in activated synoviocytes (**Fig. S7B**). The higher internalisation of psomes by activated cells possibly occurs due to the high binding affinity of the PMPC moiety to overexpressed SRB1 and CD36. In turn, these surface cell receptors bestow the uptake of psomes via endocytosis and/or phagocytosis (*37–40*). In addition, *in vitro* cytotoxicity studies assessed whether such increased uptake of psomes by activated macrophages and synoviocytes might affect cell viability. To this end, we carried out a 24-hour MTT assay confirming the biocompatibility of psomes, as both cell types did not show any considerable sign of cytotoxicity for concentrations up to 0.6 mg/mL (**Fig. S6A**). The loaded drug did not affect cell viability after 24 hours incubation time (up to 20 μg/mL; **Fig. S6B-C**). Only the free (unloaded) MTX treatment displayed a concentration-dependent cytotoxicity, suggested by the sharp decrease on cell viability at the highest concentration (**Fig. S6B-C**).

**Fig. 2.**
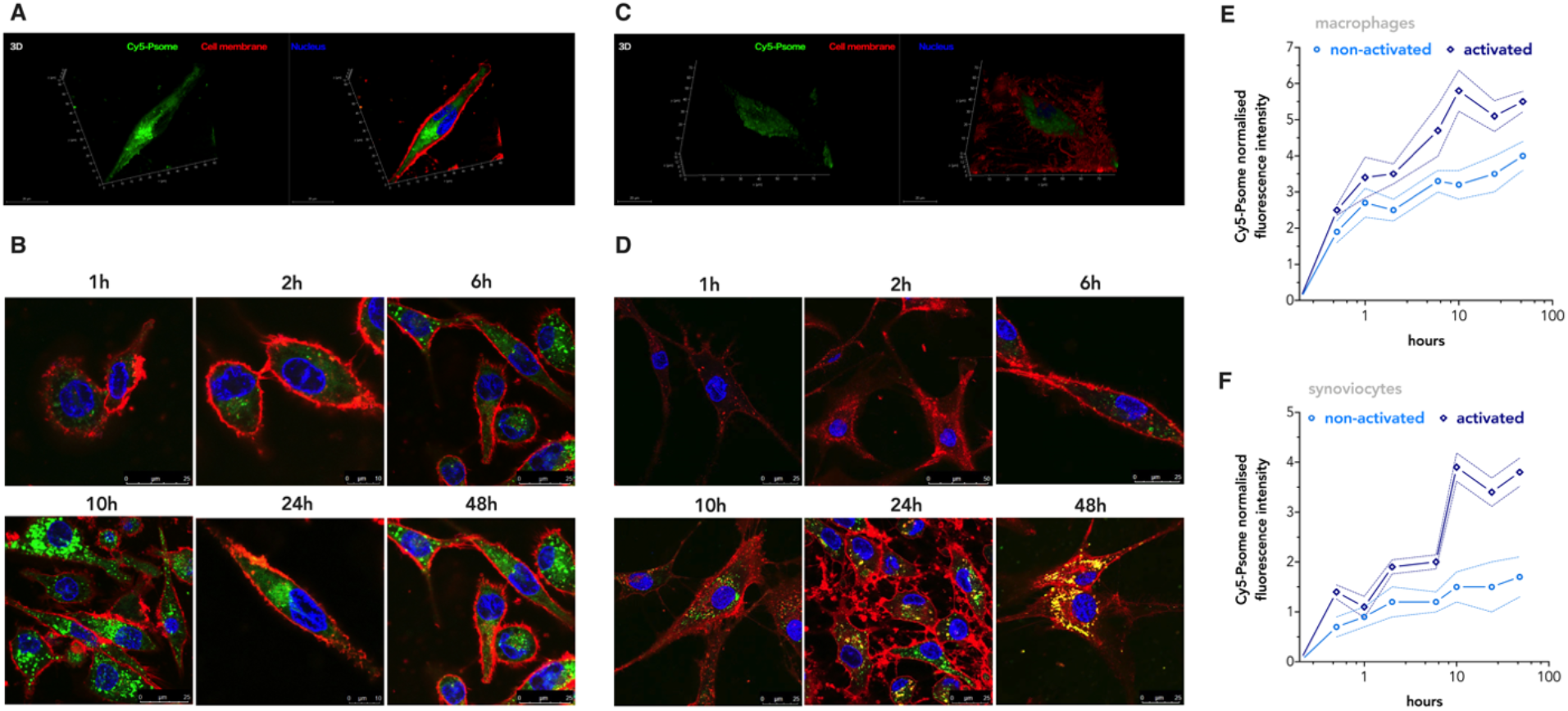
(**A**) Plasma concentration-time profile of Cy7-psomes after i.p. injection in AIA Wistar rats. (**B**) Pharmacokinetic parameters – terminal half-life (t_1/2_), elimination rate (K_e_), area under the curve (AUC) – regarding the plasma concentration-time profiles in healthy and AIA Wistar rats. (**C**) The injected dose (%ID) of Cy7-psomes per gram of organ tissue weight after 24 hours i.p. injection in healthy and AIA Wistar rats. (**D**) IVIS images of the right hind (top) and front (bottom) paws of healthy and AIA rats. (**E**) Fold change fluorescence normalised to the vehicle-treated control group. Data express the mean ± SEM (n=3 per experimental group). The differences relative to healthy group were statistically significant for *p<0.05, **p<0.01, ***p<0.001 and ****p<0.0001.

### Psomes in vitro anti-inflammatory efficacy

*In vitro* cellular studies were carried out on activated macrophages and synoviocytes to evaluate the anti-inflammatory effect of MTX-loaded psomes in comparison concerning free drug treatment. To this end, we used CLSM to investigate the cellular localisation of the transcription factor NF-κB. Macrophages and synoviocytes stimulation with LPS and TNF activates the NF-κB inflammatory signalling pathway. CLSM imaging analyses confirmed that upon celular activation, NF-κB translocates from the cytosol to the nucleus, resulting in a marked fluorescence co-localisation (quantified by the Pearson coefficient in **Fig. 3A** and **C**). Quantitative co-localisation imaging analyses demonstrated a 2-fold increase of the NF-κB nuclear translocation on both activated cell types (p<0.0001 comparing with non-activated cells; **Fig. 3B**). NF-κB is a key regulator of gene transcription in RA pathogenesis, with an important role in the production of pro-inflammatory mediators (*6, 7, 12*). Thereby, inhibiting NF-κB translocation is a crucial step to regulate the inflammatory process. CLSM imaging co-localisation analyses on both activated cell types demonstrated that a 24-hours treatment with MTX-loaded psomes significantly reduced NF-κB translocation to the nucleus (p<0.0001 compared to activated control; **Fig. 3B**). Nonetheless, both activated macrophages and synoviocytes treated with free MTX also showed a decrease nuclear translocation of NF-κB by 20 and 25%, respectively (p<0.0001 and p<0.001 compared to activated co ntrol; **Fig. 3B**).

**Fig. 3.**
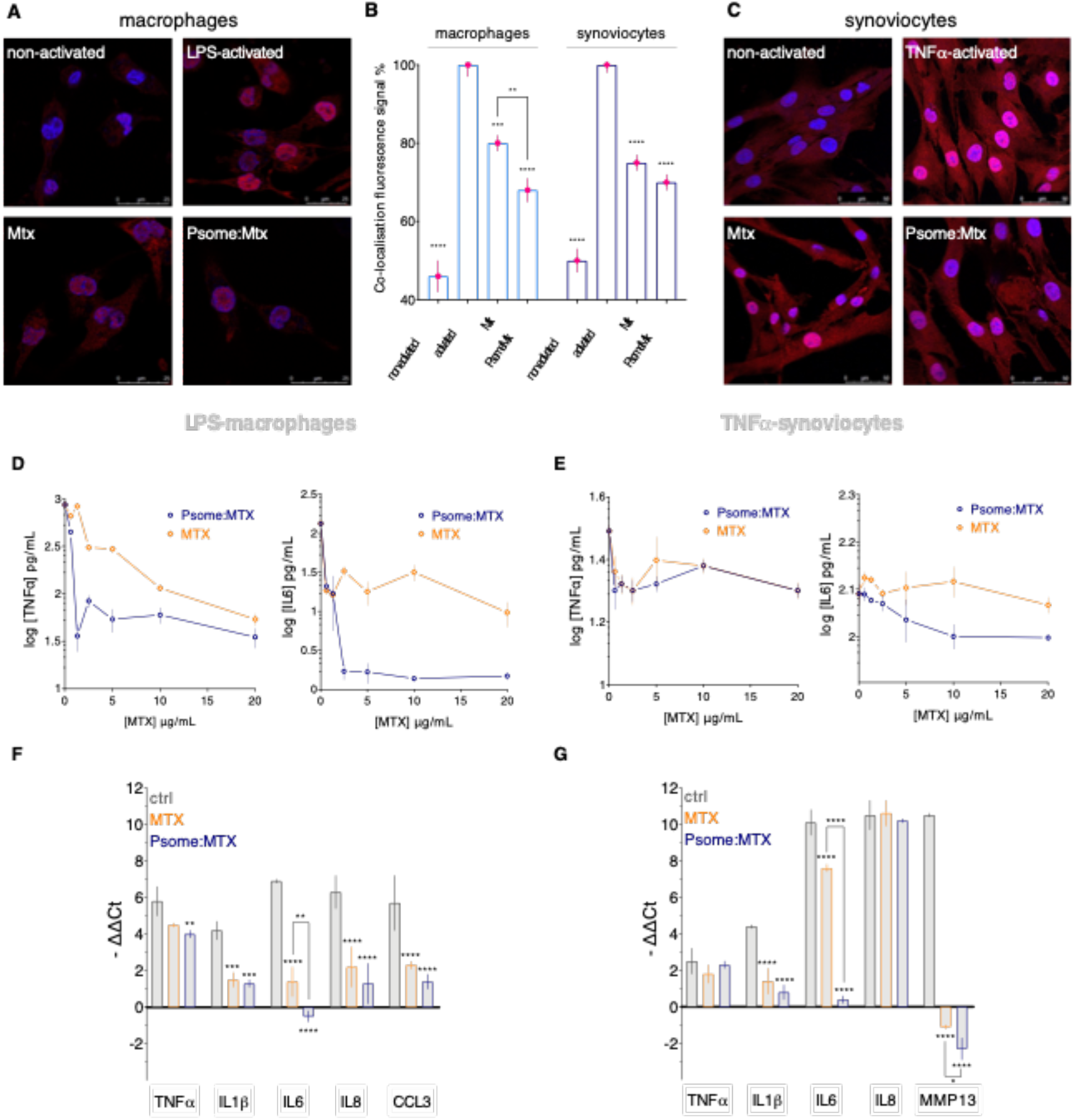
CLSM representative images of NFκB (red fluorescence intensity signal) translocation from cytoplasm to the nucleus (blue fluorescence intensity signal) on **(A)**macrophages and **(C)**synoviocytes. **(B)**Co-localisation fluorescence intensity signal (pink) analysis. Data express as mean ± SEM (5 images for n=2). ELISA analysis on the TNFα and IL6 protein secretion levels in (**D**) LPS-activated macrophages and (**E**) TNFα-activated synoviocytes. Data express as mean ± SD (n=3). RT-qPCR on the gene expression levels in (**F**) LPS-activated macrophages and (**G**) TNFα-activated synoviocytes. Data express as mean ± SD (n=3). The statistically significant differences relative to non-treated activated control for *p<0.05, **p<0.01, ***p<0.001 and ****p<0.0001.

Having demonstrated that both treatments inhibited the activation of the NF-κB signalling pathway, we evaluated the secretion of the key pro-inflammatory cytokine TNFα and IL6, which are involved in the synovial inflammatory response progression. We used ELISA to quantifie their expression on activated macrophages and synoviocytes after 24-hours treatment with increasing concentrations of either unloaded or MTX- loaded psomes. Both treatments reduced cytokine secretion levels in a concentration-dependent manner on activated macrophages (**Fig. 3D**). MTX-loaded psomes showed a strong effect in reducing the secretion level of both cytokines, which could be possibly due to their enhanced internalisation by LPS-activated macrophages (**Fig. 3D**). This suggests that psomes enable the intracellular availability of MTX. In fact, even low concentrations of loaded MTX (2.5 μg/mL) reduced TNFα levels by 1.5-fold compared with free drug treatment (**Fig. 3D**). IL6 concentration was reduced to almost undetectable levels when activated macrophages were treated with MTX-loaded psomes at concentrations higher than 2.5 μg/mL. Equal concentrations of the unloaded drug resulted in ~5-fold higher levels of the same cytokine (**Fig. 3D**). ELISA assays over activated synoviocytes also revealed that both treatments have the same effect on the secretion of TNFα, decreasing its concentration by nearly 1-fold compared to the non-treated control (*i.e.* 0 μg/mL of MTX; **Fig. 3E**). IL6 secretion levels decreased in a concentration-dependent down to 10 μg/mL in activated synoviocytes treated with MTX-loaded psomes, oppositely to free drug treatment that were maintained (**Fig. 3E**). We then evaluated the expression of inflammation-related genes on both activated synoviocytes and macrophages by RT-qPCR after 24-hours treatment with either unloaded or MTX- loaded psomes. Results confirmed that activation of macrophages and synoviocytes induces the up-regulation of all tested pro-inflammatory genes (control; **Fig. 3F-G**). Conversely, macrophages treatment with both unloaded and loaded MTX significantly reduced the expression of all the cytokines and chemokines tested (**Fig. 3F**). Both treatments also significantly reduced IL1β, IL6 and MMP13 gene expression levels on synoviocytes (**Fig. 3G**). IL6 gene expression was significantly decreased on both cell types when treated with MTX-loaded psomes (p<0.01 and p<0.0001 comparing with the free drug-treated macrophages and synoviocytes, respectively; **Fig. 3F-G**). These results demonstrate MTX-loaded psomes efficacy in reducing *in vitro* inflammation.

#### Psomes in vivo therapeutic efficacy in AIA rats

The AIA animal model has a rapid and severe disease progression following the administration of a foreign antigen of mycobacterial origin (*45, 46*). The joint swelling of the paws is indicative of RA progression and severity. At day 4 post disease induction, we observed that all hind paws of AIA Wistar rats developed arthritic inflammatory signs as evidenced by the visible erythema and swelling of the AIA joint (**Fig. 4A**). In this animal model, all the arthritic inflammatory signs are severely increased over time, reaching the acute inflammation phase by day 13 post disease induction (*47–49*). Indeed, we observed the rapid symptomatic escalation after 10 daysdays of treatment in the AIA group of animals (which also corresponds to the day 13 post induction; **Fig. 4B**). Additionally, the maximal swelling of the paws, typical in arthritic animal models, occurs by day 19 post disease induction before reaching a plateau phase (*47–49*). Taking this into consideration and having established the accumulation of psomes to inflamed synovial joints, the therapeutic efficacy of psomes to control the progression of arthritic inflammation was evaluated for 15 days of treatment in AIA rats (which corresponds the 4^th^ and 19^th^ days post disease induction time frame). To this end, the different groups of AIA rats were daily i.p. injected with either vehicle, MTX (*i.e.,* free drug), MTX-loaded psomes or empty psomes. Then, all animals were monitored constantly for 15 days by evaluating the arthritic inflammation score, which is based on assessing the ankle swelling and body weight. We observed the continuous disease progression in AIA vehicle-treated rats by their body weight loss, the increased swelling of the joints and hence of the arthritic inflammation score (**Fig. 4B-C**). Indeed, the AIA vehicle-treated group showed a significant increase of the ankle perimeter (p<0.001 comparing with the healthy control group where the arthritic inflammation score is 0; **Fig. 4C**). Conversely, all the other treatments resulted in a decrease of inflammatory arthritic signs. We observed that AIA animals treated with either unloaded or loaded MTX exhibited a significantly lower arthritic inflammation score than the psomes treatment group, especially in the last 5 days of treatment (p<0.0001; **Fig. 4B**). The average ankle swelling of the MTX-loaded psomes and MTX-treated groups was significantly reduced, respectively, by 22% and 19% relative to the first day before treatment starts (p<0.0001 *versus* the AIA vehicle-group; **Fig. 4D**). The paw swelling in AIA rats was reduced only by 4% the psomes treatment group, but still this represents a striking difference compared with the AIA rats by the end of 15 days (p<0.0001; **Fig. 4D**). AIA rats receiving either free MTX or MTX-loaded psomes treatment resulted in the same degree of paw inflammation and swelling (**Fig. 4B-D**). Moreover, we observed that AIA animals lost 6% of their weight since the beginning of treatment, oppositely to the MTX-loaded psomes group with 11% of body growth (p<0.0001; **Fig. 4E-F**).

**Fig. 4.**
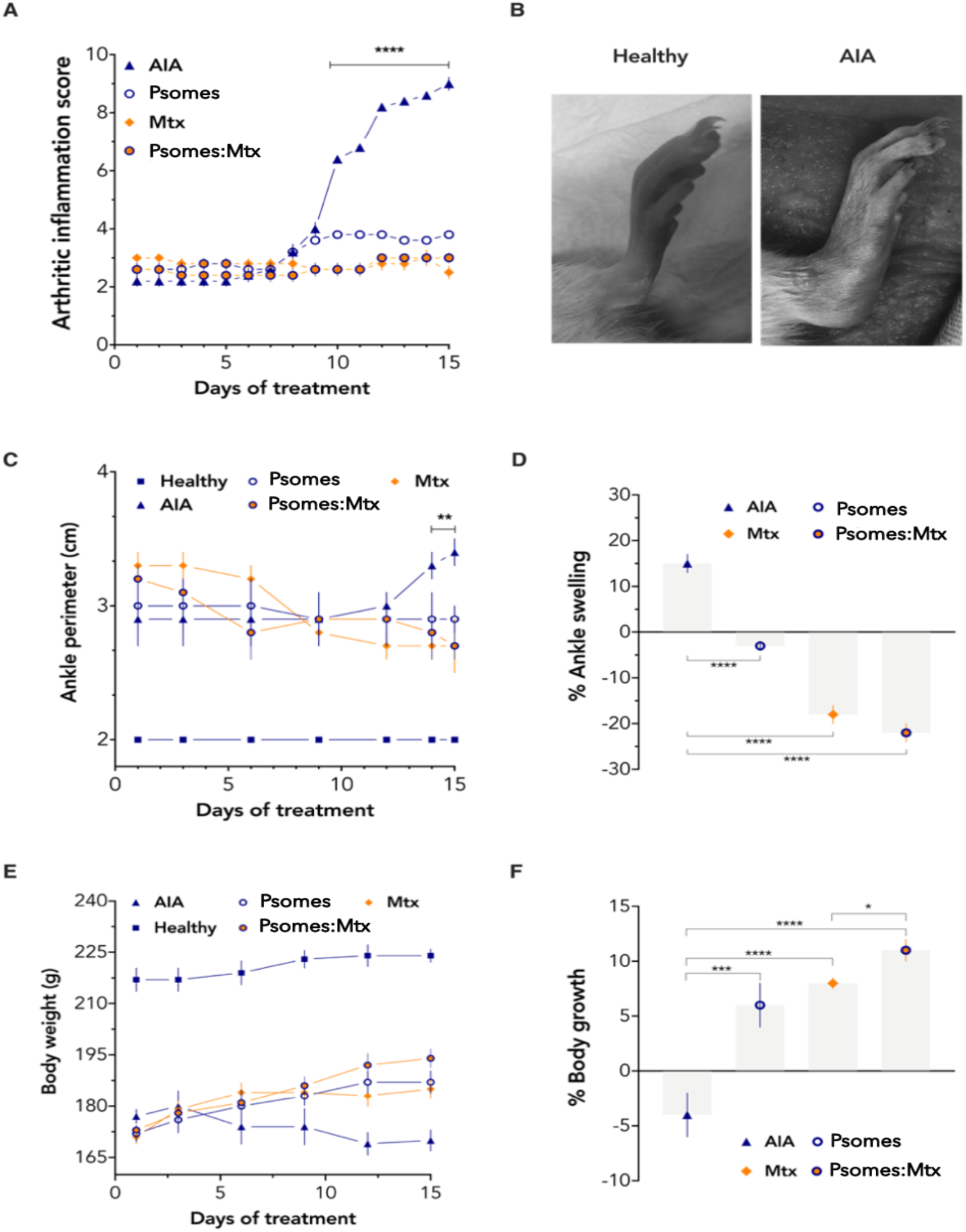
(**A**) Representative images of the left hind paw of a healthy and AIA Wistar rat. (**B**) Paws arthritic inflammation score over the 15 days period of treatment. Data defined as the mean ± SEM of the sum of the partial scores of each affected joint. (**C**) Ankle perimeter of hind paws and (**D**) variation of the ankle swelling percentage over the 15 days of treatment. (**E**) Body weight and (**F**) variation of the body growth percentage over the 15 days of treatment. Data express the mean ± SEM (n=5 per experimental group). Statistical analysis for * p ≤ 0.05, ** p ≤ 0.01, *** p ≤ 0.001 and **** p ≤ 0.0001.

The therapeutic effect of the different treatments in AIA rats joints was further evaluated using hematoxylin & eosin (H&E) histological assessment. The H&E imaging analysis of the AIA vehicle-treated animals showed signs of the early inflammatory process, such as immune cellular infiltration and proliferation, followed by formation of pannus (**Fig. 5A-C**). Histological evaluation using semi-quantitative scores was performed to identify signs of cartilage degradation and bone erosion, also revealing that both scores were significantly increased in the AIA vehicle-treated group comparing with the healthy one (p<0.0001; **Fig. 5D-E**). Conversely, H&E histopathological analyses of the AIA rat joints from either unloaded or loaded MTX treatment groups resulted in a reduction of the sub-lining layer infiltration score (**Fig. 5B**) and number of lining layer of cells score compared to the arthritic group (p<0.01; **Fig. 5C**). As a consequence of no apparent pannus formation, histological analyses of arthritic rats treated with MTX-loaded psomes showed a smooth cartilage surface and a reduced subchondral bone leukocyte infiltration (**Fig. 5E**). Consistent with these results, treatment with MTX-loaded psomes was more efficient in reducing the bone damage (p<0.1 *versus* MTX-treated group; **Fig. 5D**). Hence, negligible differences in the global disease severity score were observed in AIA rats administrated with the MTX-loaded psomes compared with the healthy rats (**Fig. 5F**). Interestingly, i.p. injection of psomes also demonstrated to have a significant therapeutic effect in the arthritic animals, particularly in preventing bone and cartilage degradation (p<0.001 and p<0.0001 *versus* AIA vehicle-treated animals respectively; **Fig. 5D-E**). Also there was no visible inflammation in the paws for all the other treatments concerning the AIA group (**Fig. 5A**). We also performed H&E histological assessment of all the livers and spleens to confirm the extent of any tissue damage or deleterious effect related to any treatment administrated. Histopathological analysis of liver sections showed the characteristic architecture with formation of lobules and normal lobular structure can be identified (**Fig. S8**), as well as the characteristic follicular structure of the spleen (**Fig. S9**). Histopathological analysis also revealed the absence of any sign of inflammation or damage in most of the livers and spleens. Apart from the livers of the MTX treated AIA rats (**Fig. S8D**), where we observed signs of fibrosis and indefinitely giant cells, possibly accompanied by mild to moderate inflammation.

**Fig. 5.**
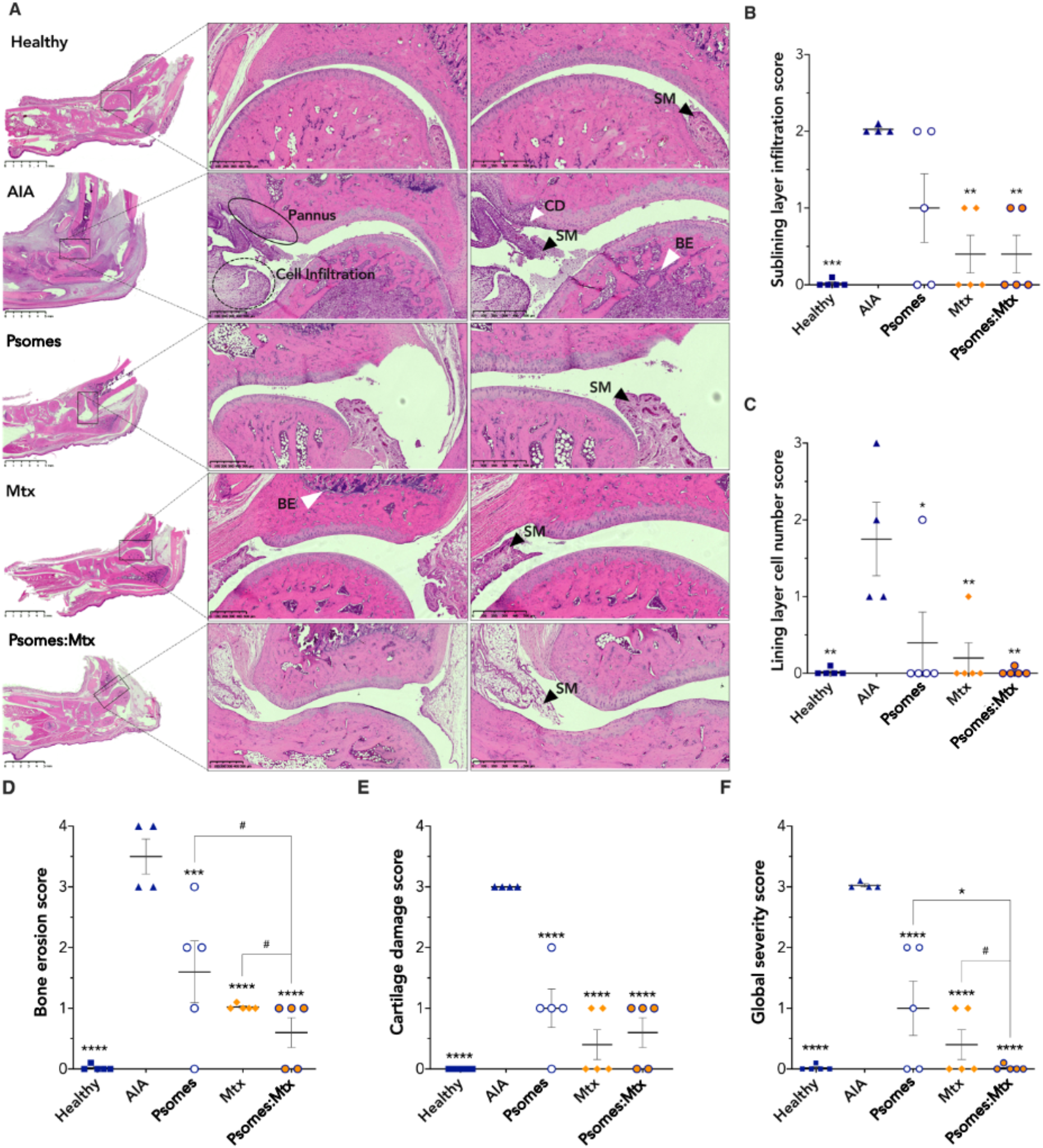
(**A**) H&E histological representative images of the left hind paw of each experimental animal group (scale bar: 500 μm; SM: synovial membrane; CD: cartilage damage; BE: bone erosion). Histological evaluation: (**B**) Sublining layer infiltration score (0 = none to diffuse infiltration, 1 = lymphoid cell aggregate, 2 = lymphoid follicles, 3 = lymphoid follicles with germinal centre formation); (**C**) Lining layer cell number score (0 = fewer than three layers, 1 = three to four layers, 2 = five to six layers, 3 = more than six layers); (**D**) Bone erosion score (0 = no erosions, 1 = minimal, 2 = mild, 3 = moderate, 4 = severe); (**E**) Cartilage damage score (0 = normal, 1 = irregular, 2 = clefts, 3 = clefts to bone); (**F**) Global severity score (0 = no signs of inflammation, 1 = mild, 2 = moderate, 3 = severe). Data express the mean ± SEM. Statistical analysis *versus* the AIA group (* p ≤ 0.05, ** p ≤ 0.01, *** p ≤ 0.001 and **** p ≤ 0.0001) and the MTX-loaded psomes treated AIA group (# p ≤ 0.1 and * p ≤ 0.05).

Additionally, to understand the efficacy of the different treatments in reducing systemic inflammation spreading in AIA rats, we investigated the levels of pro-inflammatory cytokines associated with disease activity *in vivo*. ELISA was used for the quantification of TNFα, IL1β and IL6 in the serum of all animal groups after 15 days of treatment. The vehicle-treated AIA rats exhibited high serum protein levels of TNFα, IL1β and IL6, in comparison with the healthy control group (p<0.0001, p<0.0001 and p<0.01, respectively; **Fig. 6**). Thereby, as expected, the arthritic group presented systemic inflammatory manifestations reflected by hyperplasia and synovial inflammation. In contrast, all tested inflammation-related cytokines levels decreased to a different extent in the three arthritic groups after 15 days of daily treatment. Both free MTX and MTX-loaded psomes treatments significantly decreased TNFα concentration in the serum by 5-fold (p<0.001 *versus* AIA vehicle-treated group; **Fig. 6A**). The serum levels of IL1β also decreased after treatment with both MTX and MTX-loaded psomes (p<0.001 and p<0.0001, respectively; **Fig. 6B**). Remarkably, psomes-treated AIA rats reduced the pro-inflammatory cytokine levels, particularly for TNFα and IL1β (p<0.01 and p <0.0001, respectively; **Fig. 6A-B**). Serum concentration of IL1β were similar and for both psomes treatments when compared with the healthy control group (**Fig. 6B**). We had shown in a previous *in vitro* study that PMPC-PDPA polymersomes effectively inhibit inflammatory pathways signalling and supress the production of TNFα, IL1β and IL6 by inflammation-activated macrophages (*50*). Additional analyses suggest that MTX-loaded psomes were more effective in reducing the IL1β and IL6 levels in serum than conventional MTX treatment (p<0.1 for both; **Fig. 6B-C**). Increased values of serum IL6 were also observed in the vehicle-treated AIA group, which have shown bone loss evidence in the histological analyses (**Fig. 5A, D**). By the end of 15 days, the IL6 concentration of the MTX-loaded psomes treatment group was significantly reduced to 154 ± 28 pg/mL (p<0.01 *versus* AIA vehicle-treated group; **Fig. 6C**), which was similar to the healthy vehicle-treated rats (131 ± 15 pg/mL). The *in vivo* results further support the hypothesis that psomes can effectively hinder arthritic inflammation and prevent joint destruction with negligible side-effects.

**Fig. 6.**
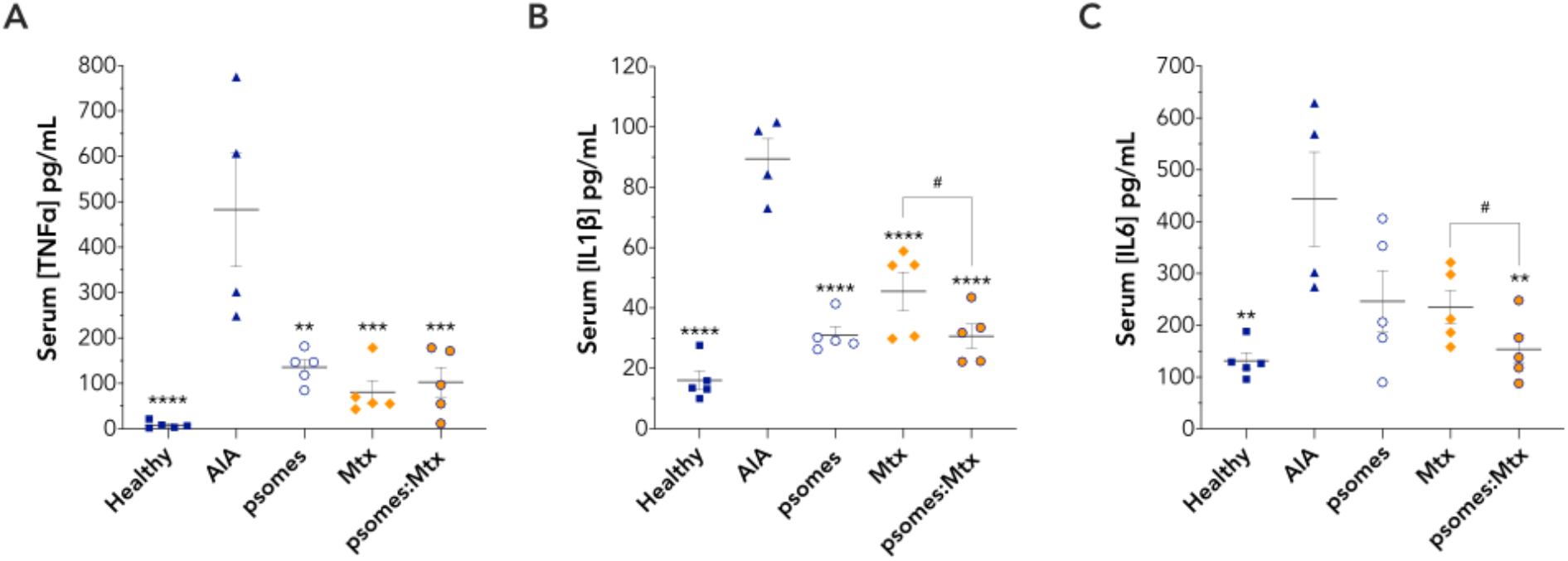
(**A**) TNFα, (**B**) IL1β and (**C**) IL6 protein levels in the serum of animals after the 15 days of treatment. Data express the mean ± SEM (n=5). Statistical analysis versus the AIA group (* p ≤ 0.05, ** p ≤ 0.01, *** p ≤ 0.001 and **** p ≤ 0.0001) and the MTX-loaded psomes treated-AIA group (# p ≤ 0.1).

## Discussion

Rheumatoid arthritis (RA) is an immune-mediated inflammatory disease associated with chronic synovial inflammation of joints (*6, 7*). At an early stage, the disease manifestations include synovitis, erythema and joint swelling (*3, 6–8, 23, 51–53*). As the disease progresses, local synovial cellular interactions and signalling cascades of pro-inflammatory mediators result in the formation of pannus tissue (*3, 5–10, 14, 51, 52*). This, in turn, causes joint stiffness and eventually leads to the irreversible damage of cartilage and bone erosion at later disease stages (*5, 6, 51, 52*). The use of MTX is the golden standard for the treatment of RA, however, it is often associated with well-known deleterious hepatotoxic side effects (*28*). In this study, we present a promising pH-responsive nanotherapy – PMPC-PDPA polymersomes(“psomes”) – with proven therapeutic potential to control the disease inflammatory activity and hinder arthritis progression, while decreasing off-target systemic exposure. Starting with *in vivo* biodistribution studies, we demonstrate the effectiveness of the nanometric size and spherical morphology of psomes to facilitate their passive accumulation within inflamed joints of AIA rats. This is most likely due to the prominent angiogenesis and enhanced vascularisation of inflamed synovial tissues and their enhanced permeation and retention effect towards nanoparticles (*9, 10, 15–18, 37, 54–57*). Additional pharmacokinetics analyses demonstrated the robust properties and stability of psomes *in vivo* upon i.p. injection. This is ascribed to the biocompatible and to the psomes hydrophilic building block, which also hinders protein fouling in the bloodstream and avoids a mononuclear phagocyte system-based clearance of the psomes (*33–36*). The accumulation of psomes in the liver and spleen of rats is, most likely, due to the fact that most i.p. injected drugs enter the portal vein and pass through the liver before entering the bloodstream. Also, these organs represent the main routes of excretion for nanomedicines (*58, 59*). More importantly, liver and spleen histopathological analyses confirm that any of the psomes treatments did not cause any tissue damage or hepatotoxic side-effects.

*In vivo* biodistribution studies are indicative of psomes ability to improve the on-site bioavailability and activity of loaded MTX within synovial disease-inflamed tissues of joints. Here, macrophages and synoviocytes found in the synovium of joints are key targets for MTX delivery, particularly due to their important role in RA pathogenesis (*6–8, 11, 12, 14*). The activation of the macrophages inflammatory phenotype induces the secretion of key pro-inflammatory mediators since the early disease stages, which are involved in the maintenance and perpetuation of the synovial inflammatory response (*6, 7, 12*). On the other hand, synoviocytes take on a more aggressive and invasive phenotype that drives the joint destruction process (*6, 11, 13–15*). With the *in vitro* cellular uptake studies, we demonstrated the psomes ability to rapidly accumulate within activated macrophages and synoviocytes without affecting their viability. We also show that this enhanced internalisation is not only ascribed to the nanometric size of psomes (*56, 60*), but mostly to their building block inherent binding affinity to the family of class B scavenger receptors, including type B1 and B3 overexpressed on these cells (*33–36*).

Upon cellular internalisation, *in vitro* drug release study confirms the acidic pH responsiveness-trigger mechanism of psomes hydrophobic building block to bestow their rapid disassemble along the endocytic pathway (*33–36, 50, 54, 57, 61–65*). This enables an efficient delivery of the loaded MTX within endosomes (pH~6 to 5) and eventual release into the cytosol, as a result of the osmotic shock in the endosome and consequent membrane poration (*37, 57, 61, 64*). In addition, this mechanism can be further exploited as a targeting strategy for RA, as the acidic environment within the synovium leads to a pH drop, thus allowing the release of loaded MTX in the target inflamed synovial tissues (*18, 66, 67*).

Further *in vitro* cellular studies gave insight into the activation of macrophages and synoviocytes as a result of the NF-κB nuclear translocation signalling and up-regulation of pro-inflammatory genes involved in the progression of chronic synovial inflammation. Additionally, we demonstrate the anti-inflammatory potential of MTX-loaded psomes in inhibiting the NF-κB signalling pathway and in modulating the expression of pro-inflammatory cytokines and chemokines. Among these, IL6 mediates both innate and adaptive immune system activation, together with TNFα and IL1β, since the early acute inflammatory state of the disease (*6, 7, 12, 68, 69*). Therefore, down-regulation of IL6 and IL1β, observed on both activated macrophages and synoviocytes, is crucial to prove psomes treatment anti-inflammatory efficacy. Moreover, the immunosuppressive effect of MTX-loaded psomes was further demonstrated by the reduced expression levels of IL8 and CCL3 genes on activated macrophages. These chemokines are involved in the acute inflammation phase of RA mediating the recruitment and activation of immune cells to the synovium (*6, 7, 12, 70*). Further confirmation of the potential efficacy of MTX-loaded psomes in shutting down the joint destructive process in RA results from the down-regulation of MMP13 gene expression on activated synoviocytes. As synoviocytes activation induces the production of these metalloproteases responsible for the cartilage damage (*6, 7, 12–15*). Also, inactivation of the NF-κB nuclear translocation on activated synoviocytes might have an impact on bone erosion because RANK (which induces osteoclast activation) is regulated by NFκB-dependent signalling (*6, 7, 11–15*). Hereby, we prove *in vitro* that MTX-loaded psomes promote inflammation resolution at the molecular level.

As the disease progresses, synovial hyperplasia and chronic inflammation eventually lead to the destruction of joints (*11, 51, 52*). Psomes were evaluated for their anti-inflammatory and anti-arthritic therapeutic efficacy *in vivo* by assessing the arthritic inflammation signs and the histopathological features of joints of AIA-treated rats. *In vivo* studies revealed the therapeutic potential of MTX-loaded psomes in the abrogation of disease progression and severity. Clinical arthritic inflammation signs and histopathological analyses on AIA rats treated with MTX-loaded psomes resulted in the suppression of synovial inflammation and also in the complete abrogation of joint synovial hyperplasia, and hence prevented cartilage and bone damages. The enhanced on-site bioavailability of MTX is fundamental for its therapeutic efficacy. Hence, the anti-rheumatic and -inflammatory therapeutic effect of psomes mostly results from their facilitated accumulation within synovial tissues of inflamed joints. However, we also show that MTX treatment had similar beneficial therapeutic effect in the inhibition of the inflammatory joint signs. The timing and dosing of MTX administrated are also key parameters for treatment efficacy, due to the drug’s rapid clearance. It should be noted that the same dose of either unloaded or loaded MTX (0.223 mg/kg/day) was used.

We demonstrated the efficacy of MTX-loaded psomes to inhibit systemic inflammation spreading in arthritic animals by analysing serum levels of pro-inflammatory cytokines. In RA, synovial inflammation progresses through interacting cascades of pro-inflammatory cytokines. TNFα, IL1β and IL6 are reported to be inherently associated with systemic and local inflammations in patients with RA (*6, 7, 11, 12*). TNFα and IL1β are believed to play an important role since the early phase of the disease as they directly stimulate the infiltration of leucocytes, neutrophils and macrophages into the synovium (*6, 7, 11, 12*). As mentioned before, IL6 is involved in the perpetuation of synovial inflammation together with IL1β and TNFα, but it plays also a key role in the progressive damage of joints (*6, 7, 11, 12, 68, 69*). IL6 activates the synoviocytes to release metalloproteases and reactive oxygen species, leading to the destruction of the cartilage tissue (*6, 7, 11, 12, 14, 68*). This cytokine is also involved in osteoclast differentiation and activation, which are mostly responsible for bone erosion (*6, 7, 11, 12, 14, 68*). Thereby, reduced levels of these inflammation-mediated cytokines confirm the potential of psomes to control the progression of systemic inflammation, as well as to hamper the disease destructive process. Additionally, the safety and biocompatibility of psomes in prolonged administration must be guarantee, as they should not cause any cytotoxic, inflammatory, or immunogenic effects. To this respect, we showed the biocompatibility of psomes *in vivo,* as no subacute systemic toxicity was observed after 15 days of daily treatment. Remarkably, we show that psomes alone is also effective in supressing joint and systemic inflammatory signs in AIA model of arthritis. Previous reports ascribed the anti-inflammatory action of psomes to their enhanced internalisation through the cell scavenger receptor class B type 1, which, in turn, is involved in the regulation of intracellular inflammatory pathways (*44, 50, 71*). We show that polymersomes made of PMPC-PDPA can indeed target macrophages and deliver drugs intracellularly, making them an ideal candidate to target and treat RA. Psomes accumulation in the inflamed joints of AIA rats, followed by selective delivery of MTX within inflamed macrophages and synoviocytes, enhanced its on-site therapeutic activity. Psomes loaded with MTX exhibit an effective anti-arthritic and anti-inflammatory therapeutic effect with significant amelioration of synovial inflammation and complete abrogation of arthritis progression and severity *in vivo*. In particular, the modulation of pro-inflammatory expression profiles seems to be crucial for shutting down synovial inflammation *in vitro*, by acting at the molecular level of the pathophysiological cascade. Thereby, psomes improve the therapeutic efficacy of loaded MTX in resolving synovial inflammation, while minimizing its deleterious off-site effects that compromise the effectiveness of conventional RA treatment.

## Materials and Methods

### Preparation and characterisation of psomes

Polymersomes formulations are made of the highly biocompatible hydrophilic – poly(2-methacryloyloxyethyl phosphorylcholine) (PMPC) – and hydrophobic poly(2-(diisopropylamino)ethyl methacrylate) (PDPA) polymer blocks designed to self-assemble into vesicles in aqueous conditions at pH above 6.2 (the PDPA pKa) (*50, 61–63*). Formulations of PMPC-PDPA polymersomes (*i.e.* psomes) were then prepared using a previously reported pH-switch method with some modifications (*65, 72*). Briefly, under sterile conditions, PMPC_25_-PDPA_68_ amphiphilic polymer, previously synthesised by atom-transfer radical polymerization (*73*), was dissolved in acidic phosphate-buffered saline (PBS 0.1 M at pH 2.0, Sigma-Aldrich) up to a concentration of 10 mg/mL. The pH-driven self-assembly process was controlled by increasing the solution pH from 2.0 to approximately neutral pH 7.4, through an injection system (2 μL/minute) of sodium hydroxide (NaOH 0.5 M at pH~14). Formulations of psomes loaded with methotrexate (MTX, C_20_H_22_N_8_O_5_, MW 454.44, λ (300 nm), Sigma-Aldrich) were prepared also using the above-mentioned pH-switch method by injecting the MTX (3 mg/mL) dissolved together in the NaOH solution. Additionally, for fluorescence imaging purposes either Cyanine5- or Cyanine7-labelled psomes were prepared using the solvent-switch method as reported with some modifications (*74*). Briefly, 10% (w/w) Cy5- or Cy7-PMPC_25_-PDPA_68_ copolymer was dissolved together with the PMPC_25_-PDPA_68_ in an organic solution of 3:1 (v/v) methanol:tetrahydrofuran (Sigma-Aldrich) for a final copolymer concentration of 20 mg/mL. Then, PBS at pH 7.4 was added into the copolymer solution under constant stirring at 42°C, through needle injection system at the rate of 2 μL/minute, until a 0.6% (w/v) PBS content was reached. Finally, any remained organic solvent was removed by dialysis (3.5 kD MWCO tubing membrane, Spectrum Labs) against PBS for 2 days. Afterwards, all formulations were purified, as previously described by size exclusion chromatography, and characterised in terms of size, size distribution and morphology (*72*). Dynamic light scattering (DLS) technique was used for the hydrodynamic size measurements in the Zetasizer Nano ZS (Zen1600, Malvern Instruments), equipped with a 633 nm HeNe laser in a scattering angle of 173°.

Additionally, the morphology was accessed by transmission electronic microscopy (TEM) using a JEOL 2100 operating at 200 kV, equipped with an Orius SC2001CCD Gatan camera. Prior to DLS and TEM characterization, all samples were respectively prepared as previously reported (*50, 75*). Follow the purification process, the psomes drug loading capacity, estimated as the number of MTX molecules per polymersome, was determined using a previously reported method (*50, 62*), after measurement on the amount of PMPC_25_-PDPA_68_ copolymer and loaded MTX by high-performance liquid chromatography (HPLC, Dionex Ultimate®3000, Thermo Scientific).

The drug-polymer interaction study was performed by determining the partition coefficient (expressed as *K_p_* and log D) of MTX in a polymersomes/water system. A previously reported method of derivative spectrophotometry (*76*), was used with some modifications, to determine the *K_p_* of MTX with the polymeric membrane at pH 7.4 and 37°C. Briefly, samples with increasing concentrations of psomes (0 - 120 μM) and a fixed concentration of MTX (25 μM) at pH 7.4 were prepared and incubated for 1 hour. The absorption spectra (220 - 370 nm) were obtained at 37°C using a UV-Vis microplate spectrophotometer (Synergy HT, Biotek). The obtained experimental data were mathematically analysed using a previously reported *Kp calculator* (*76*). The second-derivative spectra were used to eliminate the light scattering effect caused by the polymersomes and improve bands resolution. Hence, by plotting the second-derivative values, at a wavelength where the scattering was eliminated (λ_min_~255 nm), as a function of the psomes molar concentrations, the *K_p_* (M-1) was determined by fitting the experimental data using a nonlinear least-squares regression method (*76*). Results on the drug interaction with the water/polymer system are expressed as log *D* (details in SM: Fig. S3).

The drug release study was performed using the dialysis method under sink conditions. As previously described (*50*), 1 mL of MTX loaded psomes or free MTX (at the same concentration of the loaded one) was filled in a cellulose ester dialysis membrane tube (3.5-5 kDa MWCO, Float-a-Lyzer G2, Spectrum Laboratories Inc.). These dialyses were carried out against 10 mL of three different outer buffer pH conditions (PBS at pH 7.4; acetate-buffered solution at pH 6.5 or 5.0) for 50 hours under continuous magnetic stirring at 37°C (RT15 power, IKA-Werke GmbH & Co. KG). At regular time points, aliquots (200 μL) were withdrawn, and the same volume of respective fresh outer buffer solution replaced to maintain the sink conditions. The quantification of permeated drug aliquots throughout the 50 hours was determined by measuring the UV absorbance of MTX at λ (300 nm) using the UV-Vis microplate spectrophotometer (Synergy HT, Biotek). Mathematical models for drug-release kinetics, including zero-order and first-order equations, Higuchi and Hixson–Crowell models, were applied to each drug release profile to evaluate the mechanism of drug release (*50, 62*). The fitting of each model was evaluated based on the correlation coefficient (r2) values (details in SM; Table S1).

### Cell culture and activation

Human leukemic monocytes (THP-1) were cultured and maintained in RPMI-1640, 2 mM L-glutamine, 25 mM Hepes (Sigma-Aldrich) supplemented with 10% (v/v) heat-inactivated fetal bovine serum (FBS, Sigma-Aldrich), 1% (v/v) penicillin-streptomycin (Sigma-Aldrich) and 0.1% (v/v) amphotericin B (Sigma-Aldrich). Human fibroblast like synoviocytes (HFLS) purchase from Sigma-Aldrich were cultured and maintained in synoviocytes growth medium (Sigma-Aldrich) supplemented with 10 % (v/v) FBS, 1% (v/v) penicillin-streptomycin, 1% (v/v) L-glutamine (Sigma-Aldrich) and 0.1% (v/v) amphotericin B. Prior to all *in vitro* cellular studies, THP-1 cells differentiation into mature macrophages-like state was induced through incubation with 10 ng/mL of phorbol 12-myristate 13-acetate (PMA, Sigma-Aldrich) for 48 hours in a humidified atmosphere, 95% air, 5% CO_2_ at 37°C (*50, 77*). Moreover, unless stated otherwise, macrophages inflammatory phenotype (*i.e.* activated) was induced with 600 ng/mL of lipopolysaccharide (LPS, Sigma-Aldrich) (*50, 77*); while, inflammatory synoviocytes were induced with 20 ng/mL of TNFα (Sigma-Aldrich) (*78*). Follow by 24 hours incubation in a humidified atmosphere, 95% air, 5% CO_2_ at 37°C.

### Cell uptake imaging

The cell uptake imaging was performed using confocal laser scanning microscopy (CLSM, Leica SP8). First, macrophages and synoviocytes were seeded at a concentration of 5⋅104 cells per glass-bottom petri dish (Ibidi) and then activated to the inflammatory state as above mentioned. Then, cells were incubated with 0.5 mg/mL of Cy5-psomes for 0.5, 1, 2, 4, 6, 12, 24 and 48 hours, in a humidified atmosphere, 95% air, 5% CO_2_ at 37°C. After each incubation time point, followed by 3 steps of DPBS washing, cells were stained for CLSM live imaging. Respectively, for nuclear and cell membrane staining, Hoechst 33342 (Sigma-Aldrich) and far-red Cell MaskTM (Life Technologies) were incubated for 10 minutes at room temperature, before visualization under CLSM. At least 10 different regions of the petri dishes were captured and analysed using the Fiji ImageJ software (version 2.0). For the quantification of Cy5-psomes within stimulated macrophages and synoviocytes, their fluorescent intensity signal was normalized relative to the nuclear intensity signal.

### Cell surface receptors quantification

The cell scavenger receptor (SR)B1 and SRB3 (commonly known as CD36) proteins expression levels in either non- or activated cells were detected by Western blotting assay. First, macrophages and synoviocytes were seeded at a concentration of 106 cells per well in 6-well plate (CytoOne) and then activated to the inflammatory state as above mentioned. After 24 hours incubation in a humidified atmosphere, 95% air, 5% CO_2_ at 37°C, cells then washed with DPBS and lysed using radioimmunoprecipitation (RIPA) buffer (containing 50 mM Tris-HCl, pH 8.0, 1% Nonidet P-40, 150 mM NaCl, 0.5% sodium deoxycholate, 0.1% sodium dodecyl sulfate, 2 mM ethylenediaminetetraacetic acid and 1 mM dithiothreitol) supplemented with protease and phosphatase inhibitor cocktails. Followed by centrifugation (12,000 g) at 4°C for 10 minutes to remove the nuclei and any insoluble cell debris. The post-nuclear extracts were collected and used as total cell lysates. The protein concentration from these lysates was then determined following the Bradford assay kit protocol (Bio-Rad). Western blotting was performed as previously described with minor modifications (*79*). Briefly, the cell lysates were first denatured in 4x Laemmli sample buffer (Bio-Rad) at 95°C for 5 minutes. Then, 10 μg of total cell lysates proteins were separated by electrophoresis on 10% SDS polyacrylamide gels (previously prepared following the Bio-Rad protocol) and transferred to a polyvinylidene difluoride (PVDF) membrane (Bio-Rad). The PVDF membranes were then blocked with 5% milk in Tris-buffered saline with 0.1% Tween-20 (TBST) for 1 hour at room temperature. For the immunodetection of SRB1 and CD36 protein expression, the PVDF membranes were first incubated, overnight at 4°C, with 1:1000 dilution of each specific primary antibody (Novus Biologicals NB400-144 and NB400-131) in 1% milk/TBST. And then, after several washes with TBST, the PVDF membranes were incubated with the secondary antibodies (1:20000) in 1% milk/TBST for 1 hour at room temperature. The signals of the goat anti-rabbit and anti-mouse IgG DyLight 800 (Invitrogen) were detected using the Odyssey CLx imaging system. The densitometry analyses were performed using Fiji ImageJ software (version 2.0), and the obtained values represent the ratio between the immunodetected protein and the glyceraldehyde 3-phosphate dehydrogenase (GAPDH, Abcam) loading control. Then, the fold change expression was determined by normalisation to the non-activated control.

### Cell viability

The thiazolyl blue tetrazolium bromide (MTT, Sigma-Aldrich) assay was used, as previously reported, to evaluate the cytotoxicity of all psomes formulations (*80*). Briefly, for MTT assay, cells were seeded at a density of 5⋅103 cell per well in 96-well plates (CytoOne). After seeding and activation, increasing concentrations of each treatment were incubated for 24 hours in a humidified atmosphere, 95% air, 5% CO_2_ at 37°C. Control wells were incubated with equivalent volumes of corresponding cell culture medium and/or a solution of 10% (v/v) dimethyl sulfoxide (DMSO, Sigma-Aldrich) in DPBS. Follow by 24 hours incubation, 0.5 mg/mL of MTT solution was added to each well, and 2 hours later the MTT solution was replaced by the same volume of DMSO per well to dissolve the formed formazan crystals. Then, the optical density of solubilized blue crystals was read by measuring UV absorbance at 590 nm and 630 nm using a UV-Vis microplate spectrophotometer (SynergyTM HT, Biotek). The cell viability was determined as the percentage of the metabolic activity of treated cells normalised to the control wells.

### NF-κB signalling imaging

Nuclear factor-κB (NF-κB) signalling imaging was performed using CLSM. Firstly, cells were seeded at a concentration of 5⋅10^4^ cells per glass-bottom petri dish (Ibidi) and activated as above mentioned. Then, M1-macrophages were incubated with 10 μg/mL of either unloaded MTX or MTX-loaded psomes for 24 hours in a humidified atmosphere, 95% air, 5% CO_2_ at 37°C. Following treatment, cells were washed with DPBS and fixed using 3.7% formaldehyde (Sigma-Aldrich) for 10 minutes at room temperature. After the fixation step, followed by DPBS washing for the membrane permeabilization step, cells were incubated with 0.2% Triton-X (Sigma-Aldrich) for a further 10 minutes at room temperature. Then, the immunostaining blocking was performed using 5% bovine serum albumin (BSA) (Sigma-Aldrich), to prevent unspecific antibody binding. After 1 hour at room temperature, cells were incubated with NFκB p65 Antibody (F-6) Alexa Fluor^®^ 647 (SantaCruz Biotechnology) 1:500 diluted in 1% BSA overnight in a humidified chamber at 4°C. The following day, cells were washed with DPBS and the nucleus was stained with Hoescht 33342 (Thermo Fisher) for 10 minutes at room temperature, before visualisation under CLSM. At least 10 different regions of the petri dishes were acquired. The NF-κB nuclear translocation imaging analysis was evaluated by co-localisation (Pierce’s coefficient values) of the NF-κB and nucleus fluorescence intensity signals using Fiji ImageJ software (version 2.0).

### Enzyme linked immunosorbent assay

The IL6 and TNFα protein levels were determined using an enzyme-linked immunosorbent assay (ELISA). Firstly, macrophages and synoviocytes were seeded at a concentration of 10^6^ cells per well in 6-well plate (CytoOne) and activated to the inflammatory state as above mentioned. Followed by 24 hours treatment with 10 μg/mL of either unloaded or loaded MTX. Supernatants of each well were then collected, and the ELISA (Invitrogen) was carried following the manufacturer protocol.

### Real-time quantitative polymerase chain reaction

Analyses on the gene expression of inflammation-related markers, including TNFα, IL1β, IL6, IL8, CCL3 and MMP13 was assessed using real-time quantitative polymerase chain reaction (RT-qPCR). Firstly, cells were seeded at a concentration of 10^6^ cells/well in 6-well plate (CytoOne) and activated as above mentioned. Followed treatment with 10 μg/mL of unloaded or loaded MTX, for 6 and 20 hours, respectively for macrophages and synoviocytes. Cells were then lysed, and the RNA was extracted following the RNeasy mini kit (Qiagen) protocol pre-installed in the QIAcube (Qiagen). The total ribonucleic acid (RNA) concentration was measured with NanoDrop spectrophotometer (ThermoScientific). Complementary deoxyribonucleic acid (cDNA) was synthesised from every 1 μg of total mRNA in 20 μL volume with QuantiTect Reverse Transcription Kit (Qiagen) according to the manufacture protocol. This procedure provided a fast and efficient cDNA synthesis with integrated removal of genomic DNA contamination.

Briefly, the sample of RNA is incubated at 42°C for 2 minutes to effectively remove containing genomic DNA, then the reaction occurred for another 15 minutes at 42°C and then inactivated at 95°C. RT-qPCR reaction was performed on yield cDNA synthetized from each sample using QuantiTec® Rotor-GeneTM SYBR Green RT-PCR kit (Qiagen) using the Qiagility instrument software (Qiagen). This software enables rapid and high-precision system of sample preparation for RT-qPCR analysis, providing a step-by-step guidance for automatic calculation of all primers, cDNA template and Rotor-Gene SYBR Green master mixes need for the reaction. For the RT-qPCR experiments, the ribosomal protein L13A (RPL13A) and glyceraldehyde 3-phosphate dehydrogenase (GADPH) were used as reference genes, respectively for macrophages and synoviocytes. The list of designed primers of each target gene and reference gene are detailed in SM (Table S3). Following sample preparation, the PCR mixtures are placed in the Rotor-Gene Q cycler (Qiagen) and amplification process starts using the following protocol steps: initial cycling step at 95°C during 5 minutes for the DNA polymerase activation; followed by 40 cycles of 95°C during 5 seconds for denaturation; and 60°C during 10 seconds for combined annealing and extension for all primers. RT-qPCR data analysis of folds-changes in gene expression levels normalized to the non-activated cells was determined by the −∆∆Ct method (details in SM), using cycle threshold (Ct) values acquired from the amplification curve using the Rotor-Gene Q instrumentation software (Qiagen).

### Animal experimental design

Adjuvant induced arthritis (AIA) 8-week-old female Wistar rats purchased from Charles River laboratories international (Spain) were housed in European type II standard filter top cages (Tecniplast) at the Specific Pathogen Free animal facility at the Institute of Molecular Medicine at the Faculty of Medicine, University of Lisbon. Here, the animals were individually identified and randomly housed in experimental groups of n=5, as follows: *(i)* AIA treated with MTX (0.223 mg/kg body weight by daily intraperitoneal (i.p.) administration); *(ii)* AIA treated with MTX-loaded psomes (0.223 mg/kg/day via i.p.); *(iii)* AIA treated with psomes (10 mg/kg body weight corresponding to the an equal mass of polymer injected in the *(ii)* group); the respective *(iv)* AIA and *(v)* non-arthritic healthy vehicle groups (received an equal volume of PBS). Daily i.p. injections in treated and vehicle groups started 4 days after disease induction, when the AIA Wistar rats already presented clinical signs of arthritis (*47–49*). Experiments were approved by the Animal User and Ethical Committees, at the Institute of Molecular Medicine according to Portuguese law and European recommendations.

### Pharmacokinetics and biodistribution

Animals from both healthy and arthritic vehicle groups were injected intraperitoneally (i.p.) with Cy7-psomes polymersomes (10 mg/Kg body weight) at 24 hours prior sacrifice. For pharmacokinetics analysis, blood samples were collected from the rat tail at several timepoints (0.5, 2, 4, 8 and 24 hours) and immediately processed (centrifugation 2000 rcf for 10 minutes at 4°C) for plasma separation from the blood cells. The plasma concentration of near infrared (NIR) fluorescence of Cy7-psomes was measured using a multimode microplate fluorometer (Spark, Tecan, Switzerland). For the biodistribution study, 24 hours post i.p. injection., the main organs (kidney, spleen, liver and heart) were collected and wet weight and further NIR fluorescence analysis of Cy7-psomes was accessed. Briefly, a part of the organ tissue was weight and homogenised following the protocol of the Precellys soft tissue lysing kit (CK14, Bertin Instruments, VWR, UK). The fluorescence of Cy7-psomes was measured in using a multimode microplate fluorometer (Spark, Tecan, Switzerland). Additionally, an *in vivo* imaging system (IVIS, Perkin Elmer) was used to evaluate the biodistribution of Cy7-psomes in the front and right hind paws of both healthy and AIA Wistar rats.

### Arthritic inflammatory signs

Arthritic inflammatory signs in rats’ paws and body weight were evaluated daily during the 15 days period of treatment. Inflammation scores were recorded following standard protocols and criteria established (*47–49*) by counting the score of each joint in a scale of 0 – 3, where: 0 = absence, 1 = erythema; 2 = erythema and joint swelling; 3 = deformities and functional impairment of the entire paw. The total inflammation score of each animal was defined as the sum of the partial scores of each affected joint. Additionally, the swelling of the hind paws was evaluated by measuring the ankle perimeter. In the end, all animals were then sacrificed 19 days post disease induction (15 days after treatment was started) where a maximum disease activity and severity occurs (*46*). At the sacrifice time, animals were anesthetised with pentobarbital (100 mg/kg body weight) administrated intraperitoneally. Here, the blood samples were collected by cardiac puncture and then processed (centrifugation 2000 rcf for 10 minutes at 4°C) for further serum subsequent analysis. Afterwards, the animal was perfused transcardially with PBS for vascular and organs haemoglobin release. The main organs (kidney, spleen, liver and heart) and all paws were also collected and wet weight for further analysis.

### Histopathology

Left hind paw samples were collected at the time of sacrifice and fixed immediately in 10% (v/v) neutral buffered formalin solution and then decalcified in 10% (v/v) formic acid. Samples were then dehydrated and embedded in paraffin, serially sectioned at a thickness of 4 μm and then stained with haematoxylin and eosin (H&E) for histological analysis. The histopathological evaluation (by BV) of structural changes and cellular infiltration in paw sections was performed following previously reported criteria in a blind fashion using 5 semi-quantitative scores (*47–49*): Sublining layer infiltration score (0 = none to diffuse infiltration, 1 = lymphoid cell aggregate, 2 = lymphoid follicles, 3 = lymphoid follicles with germinal center formation); Lining layer cell number score (0 = fewer than three layers, 1 = three to four layers, 2 = five to six layers, 3 = more than six layers); Bone erosion score (0 = no erosions, 1 = minimal, 2 = mild, 3 = moderate; 4 = severe); Cartilage surface (0 = normal, 1 = irregular, 2 = clefts, 3 = clefts to bone); Global severity score (0 = no signs of inflammation, 1 = mild, 2 = moderate, 3 = severe). Additionally, sections of both liver and spleen of all experimental groups were as well stained with H&E and logically analysed by an expert pathologist (SB).

### Serum inflammation-related cytokines

Serum levels of TNFα, IL1β and IL6 were quantified using a specific rat enzyme-linked immunosorbent assay (ELISA, Invitrogen, Thermo Scientific) following the manufacturer protocol.

### Data statistical analysis

Statistical analyses were performed using GraphPad Prism (version 8.2.1). Differences between groups were assessed by one-way or two-way ANOVA with Tukey multiple comparison test. The differences were statistically significant when * p < 0.05, ** p < 0.01, *** p < 0.001 and **** p < 0.0001.

## Acknowledgments

V.M.G. thanks the received financial support from FCT (Fundação para a Ciência e Tencnologia) through the FCT PhD Programmes and by POCH (Programa Operacional Potencial Humano), specifically by the Doctoral Programme on Cellular and Molecular Biotechnology Applied to Health Sciences for the grant (PD/BD/128388/2017). V.M.G. is also grateful to Diana Matias for her expertise and technical assistance with western blotting assay. C.N. also thanks FCT for the Investigator grant (IF/00293/2015) and respective exploratory project. GB thanks ERC for the consolidator award (CheSSTaG 769798), EPSRC Established Career Fellowship (EP/N026322/1), EPSRC/SomaNautix Healthcare Partnership EP/ R024723/1, and Children with Cancer UK for the research project (16-227).

## Author contributions

V.M.G. prepared and characterised the formulations, with A.P. and C.L. contribution on the polymer synthesis and TEM, respectively. V.M.G. and L.R., designed and performed the *in vitro* experiments. V.M.G., L.R. and B.V. designed and performed the *in vivo* experiments, with E.S. and S.B. contribution on the IVIS and histological analyses, respectively. V.M.G., L.R., B.V., and C.N. contributed to data analysis and interpretation. V.M.G. prepared the figures and wrote the manuscript under the supervision of G.B., L.R. and S.R. and A.O. G.B., S.R. and A.O. supervised the research and contributed to the experiments design. All authors contributed substantially to the revisions of the manuscript and gave final approval for publication.

## Data and materials availability

All data needed to evaluate the conclusions in the manuscript are present in the manuscript and/or the Supplementary Materials. Additional data related to this manuscript may be requested from the authors.

## Supplementary Materials

**Fig. S1.**
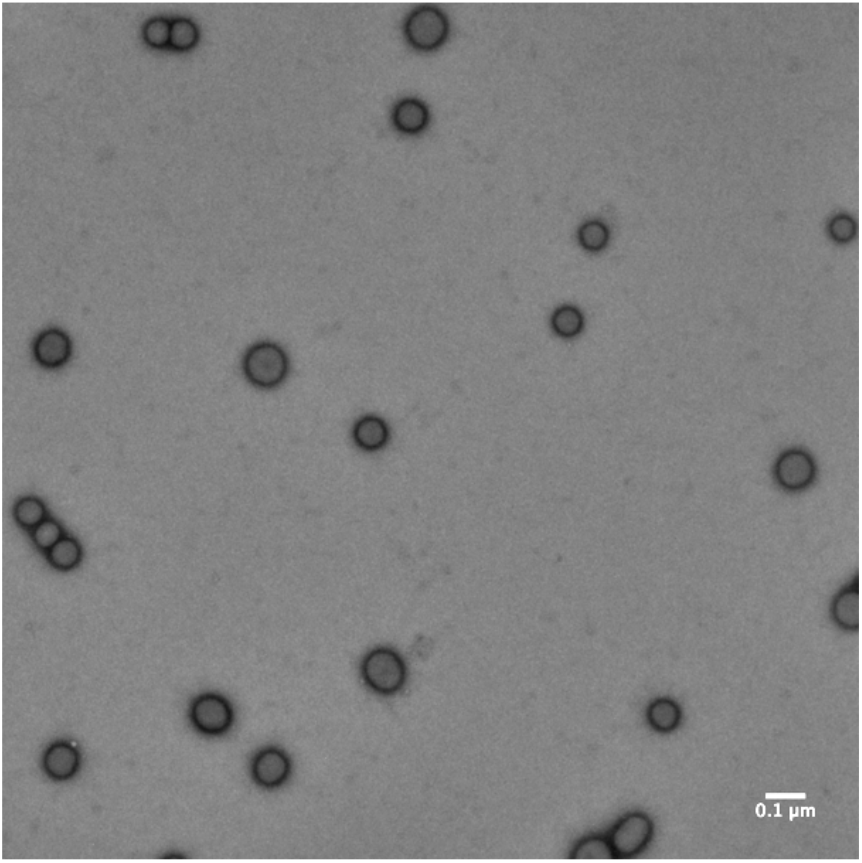
Representative TEM micrograph o f p s o m e s (*i.e.* P M P C - P D P A polymersomes) loaded with MTX produced via pH-switch (0.1 μm scale bar).

**Fig. S2.**
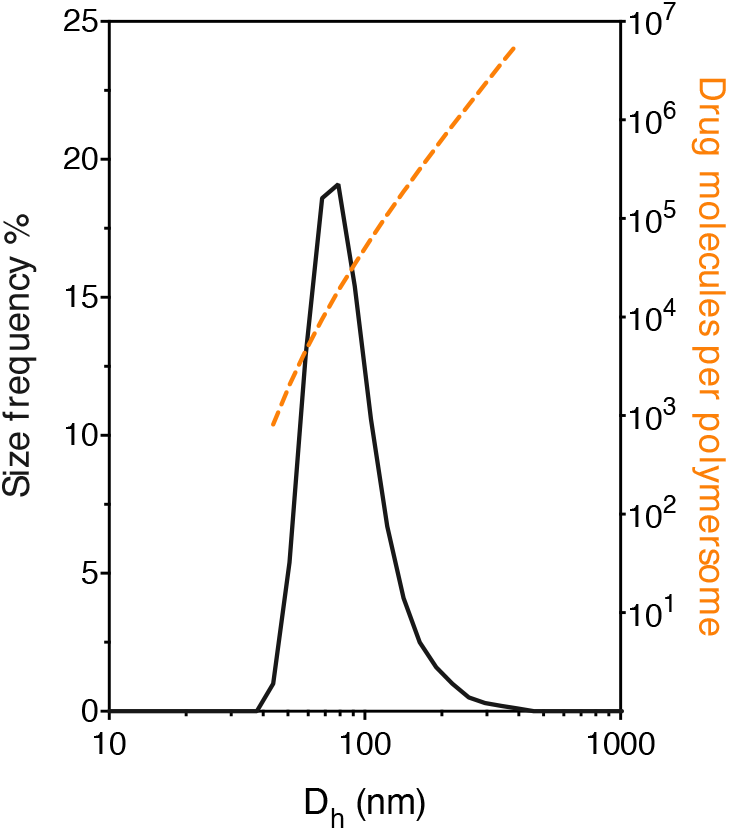
Analysis of the drug loading capacity represented as the number of MTX molecules per psome as a function of their hydrodynamic diameter (D_h_) measured by DLS.

**Fig. S3.**
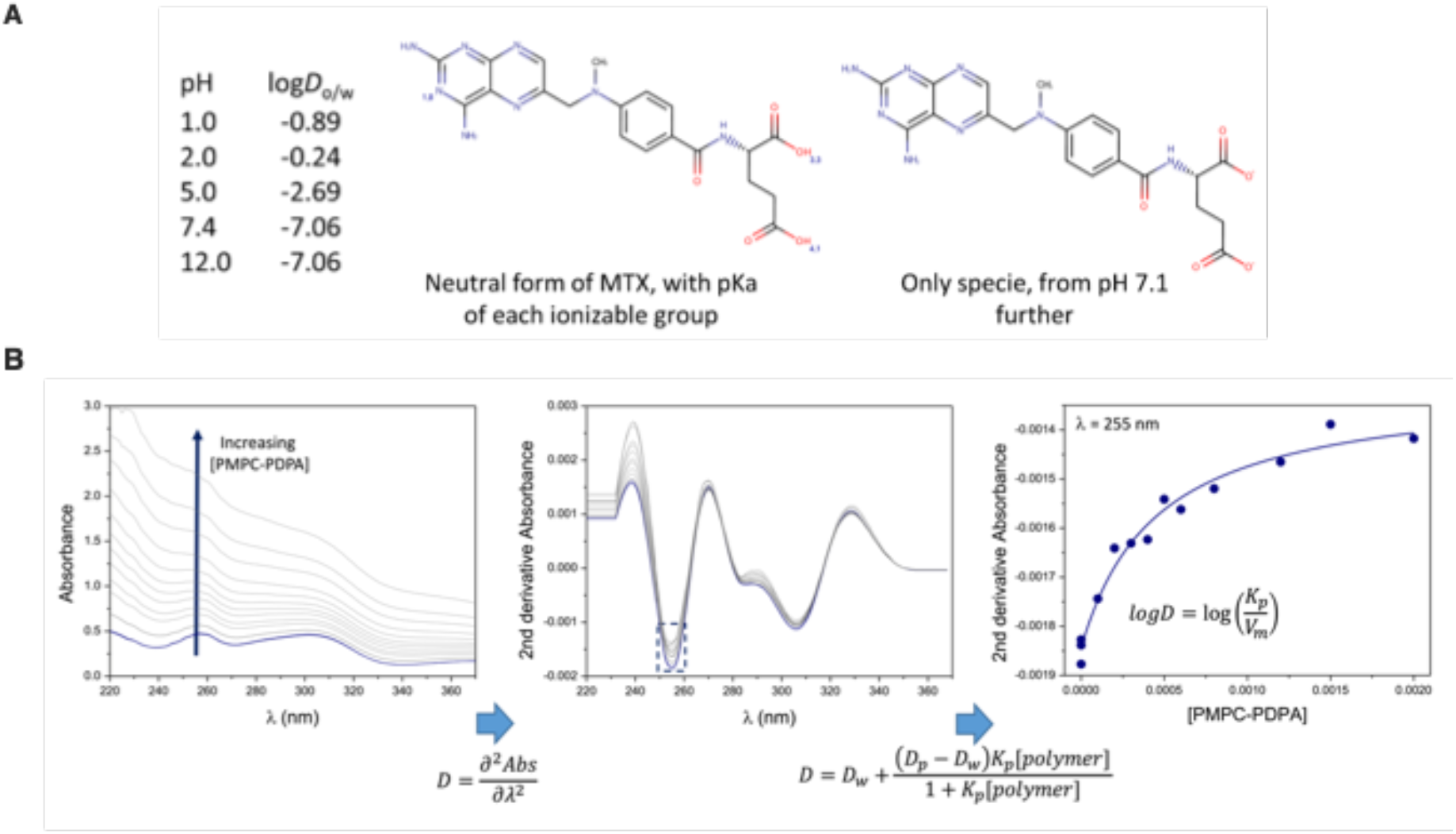
**(A)** Log *D* calculations on an octanol/water system for MTX at different pH conditions using the MarvinSketch calculator (Chemaxon). The solubility of MTX in aqueous media was found to increase as a function of the pH, due to the deprotonation of the two carboxylic acids present in the chemical structure at pH above 7.4. **(B)** Absorption spectra and second-derivative spectra of MTX (25 μM, blue lines) incubated with increasing concentrations of psomes (0 – 120 μM, grey lines) at pH 7.4 and 37°C. Graphic represents the fitting curve of the experimental second-derivative spectrophotometric data using a nonlinear least-squares regression method at wavelength 255 nm where the scattering is eliminated. The partition coefficient at pH 7.4 are then calculated by fitting the equation to experimental derivative spectrophotometric data through a nonlinear regression method, where the adjustable parameters are *D_p_* and *K_p_*. *D*, *D_w_* and *D_p_* correspond to the second derivative of total, aqueous and polymer absorbance of MTX, respectively; *K_p_* is the partition coefficient (M^−1^), *[polymer]* the psomes concentration (mol.L^−1^) and *V_φ_* the polymer molar volume (L.mol^−1^).

**Fig. S4.**
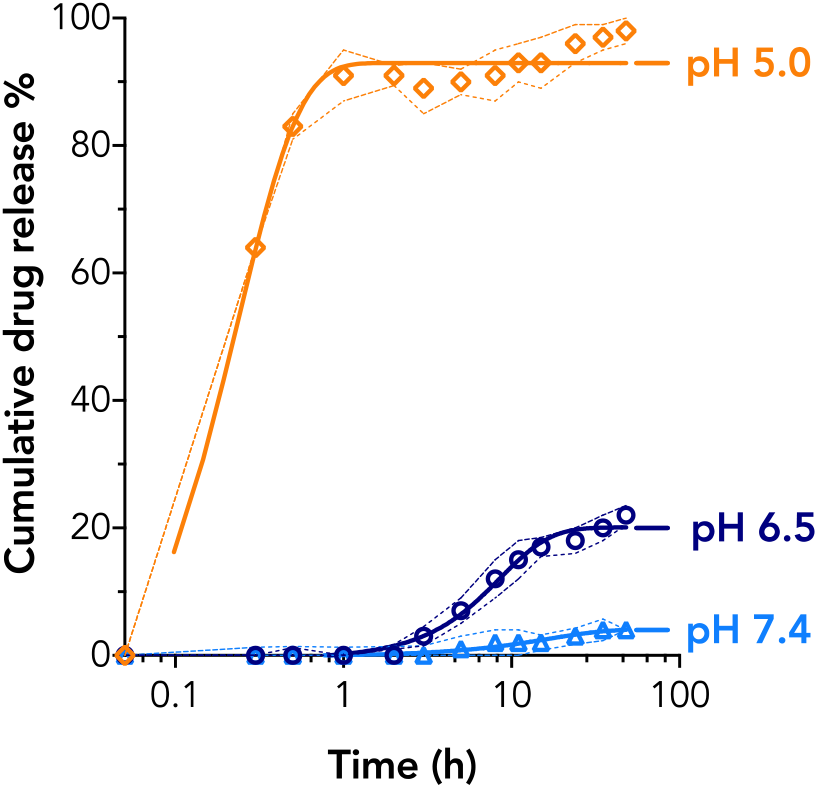
Drug release profiles of MTX from psomes polymersomes in pH 5.0, 6.5 and 7.4 buffer solutions at 37°C for 50 hours (n = 3).

**Table S1.** Correlation coefficient (r^2^) from various drug release mathematical models for each pH profile.

|  | Zero-Order | First-Order | Hixson-Crowell | Higuchi |
| --- | --- | --- | --- | --- |
| *pH 5.0* | 0.890 | 0.348 | 0.648 | 0.994 |
| *pH 6.5* | 0.957 | 0.561 | 0.708 | 0.962 |
| *pH 7.4* | 0.676 | 0.462 | 0.573 | 0.684 |

**Table S2.**
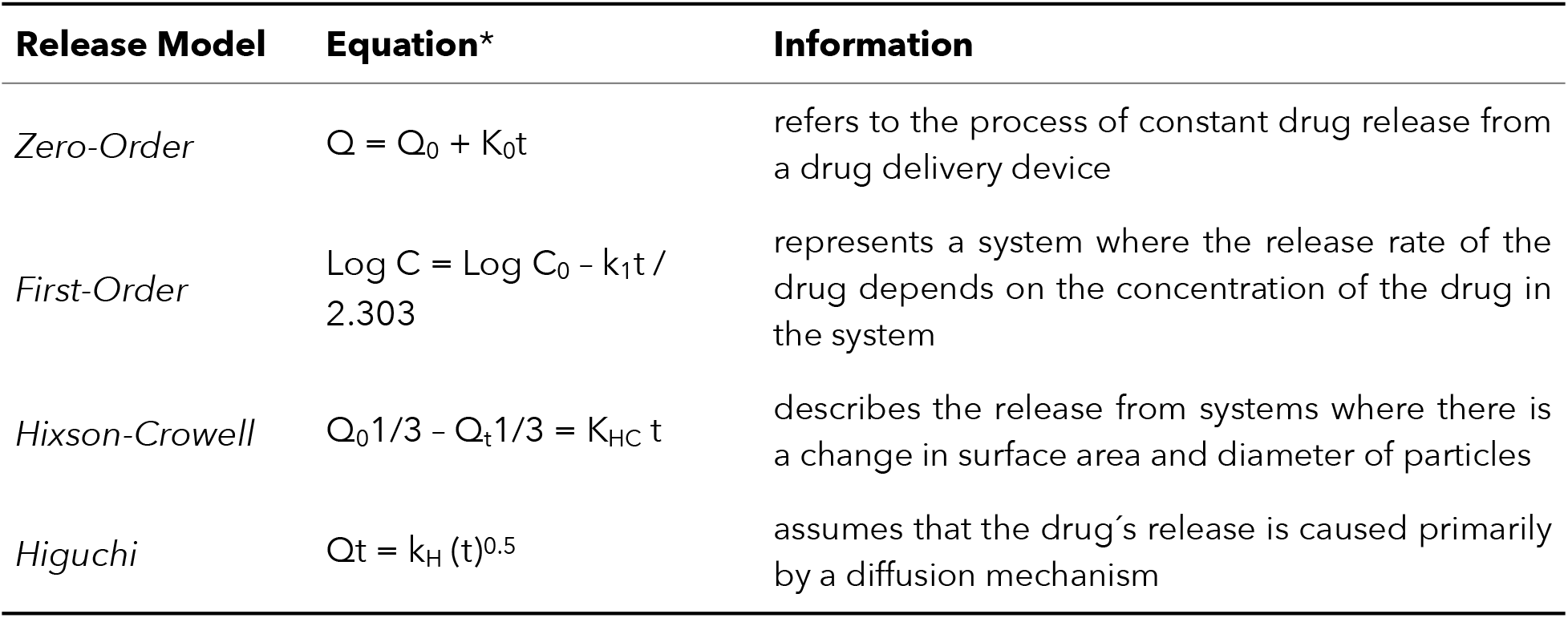
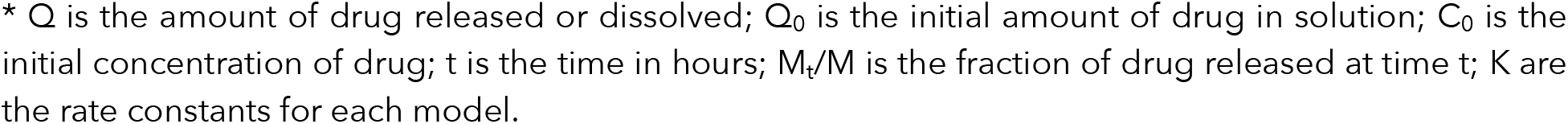
Mathematical models for drug-release kinetics.

| Release Model | Equation* | Information |
| --- | --- | --- |
| *Zero-Order* | $Q = Q_0 + K_0t$ | refers to the process of constant drug release from a drug delivery device |
| *First-Order* | $\text{Log } C = \text{Log } C_0 - k_1t / 2.303$ | represents a system where the release rate of the drug depends on the concentration of the drug in the system |
| *Hixson-Crowell* | $Q_0^{1/3} - Q_t^{1/3} = K_{HC} t$ | describes the release from systems where there is a change in surface area and diameter of particles |
| *Higuchi* | $Q_t = k_H (t)^{0.5}$ | assumes that the drug's release is caused primarily by a diffusion mechanism |

\* Q is the amount of drug released or dissolved; $Q_0$ is the initial amount of drug in solution; $C_0$ is the initial concentration of drug; t is the time in hours; $M_t/M$ is the fraction of drug released at time t; K are the rate constants for each model.

**Table S3.**
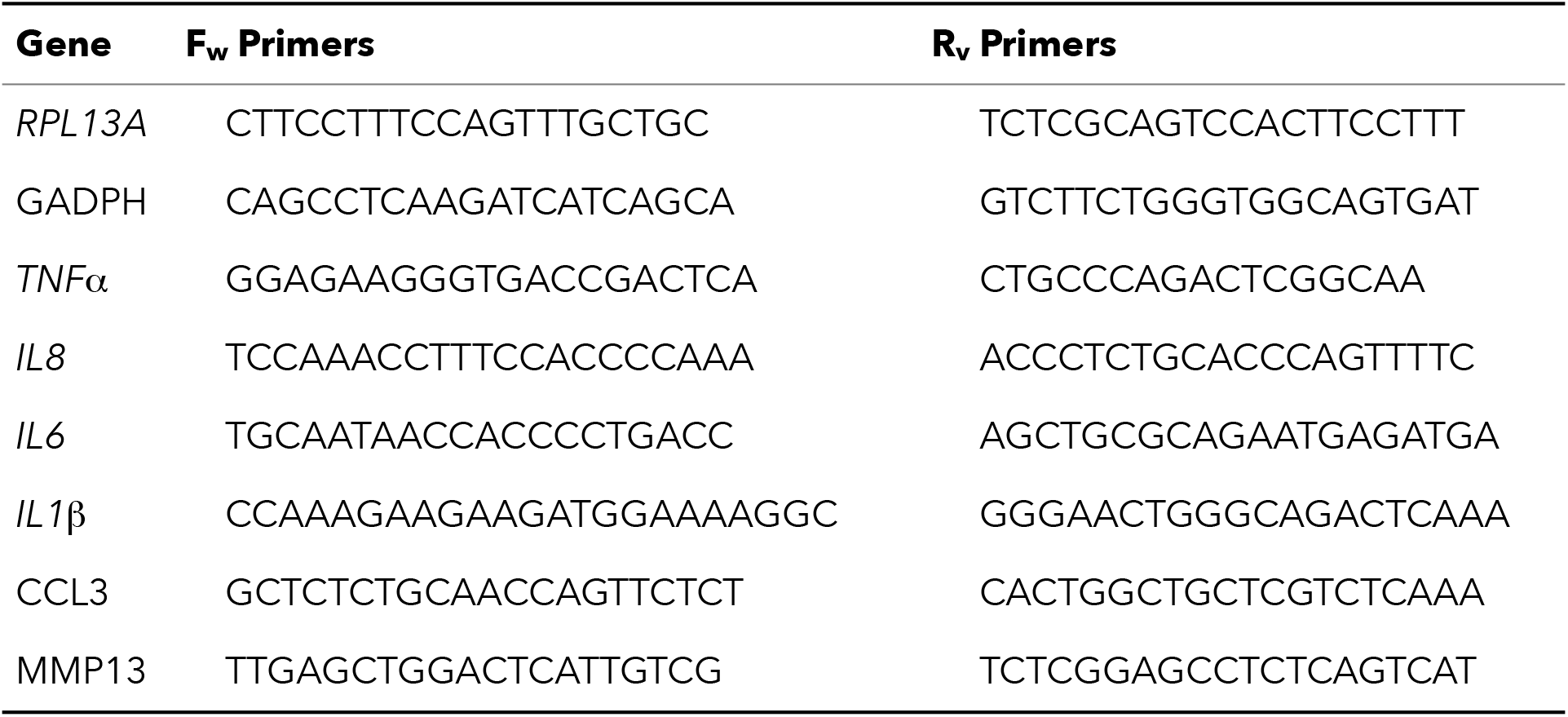
Forward (F_w_) and reverse (R_v_) gene sequences of designed primers (PRIMER-BLAS from Sigma-Aldrich) used for gene expression studies. RT-qPCR data was analysed using the comparative cycle threshold (Ct) method, also known as the ∆∆Ct method. The Ct value of each target gene was normalized to the reference gene, obtaining the ∆Ct value (eq. 1) of treatment and non-treated control. Then, the change in Ct is compared against the control to obtain the ∆∆Ct value (eq. 2). Then, the –∆∆Ct values corresponds to the folds in gene expression change of the treated compared to the non-treated group.

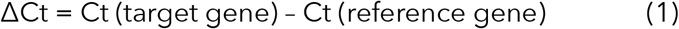

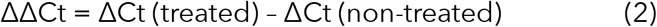

**Fig. S5.**
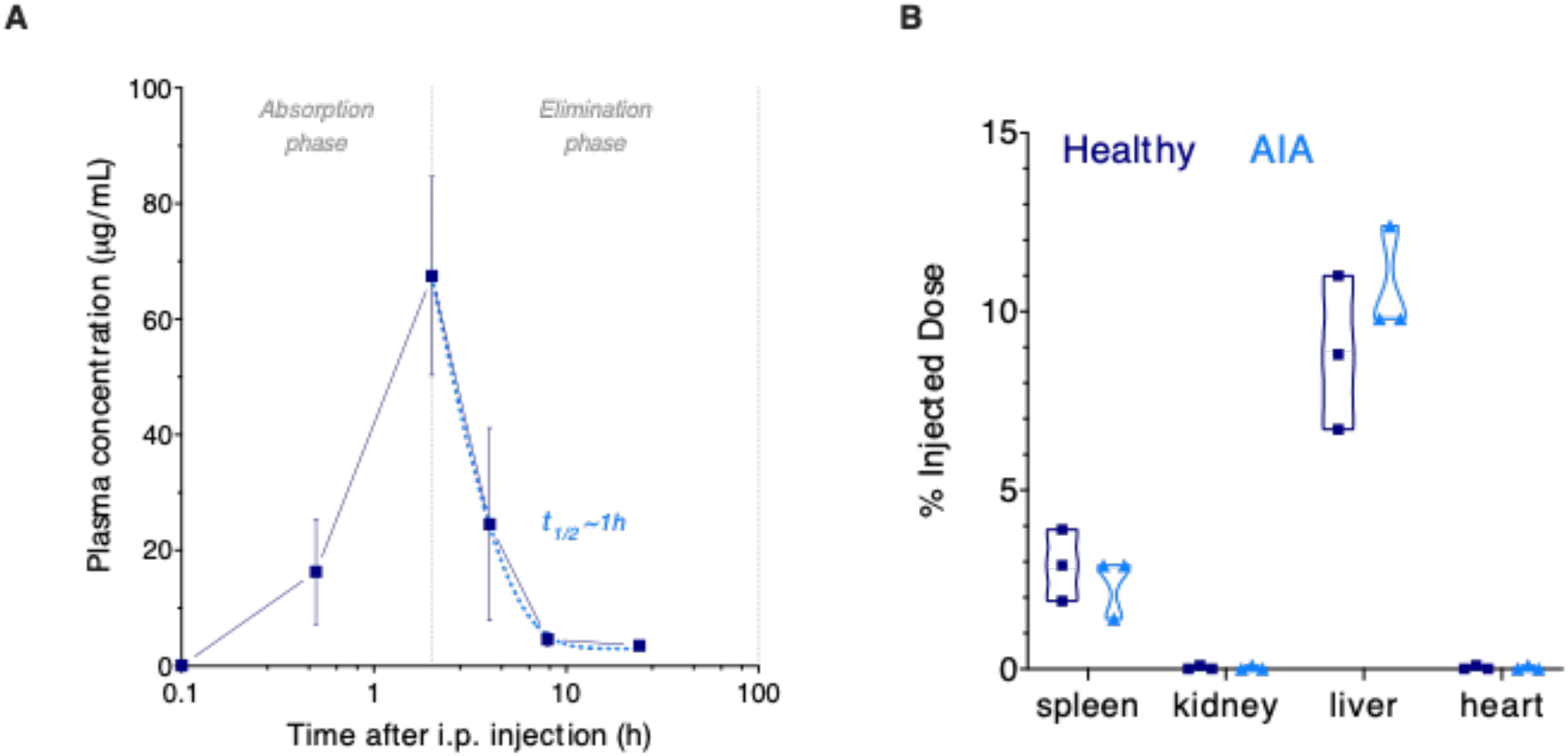
(**A**) Plasma concentration-time profile of Cy7-psomes after i.p. injection in healthy Wistar rats. Terminal half-life (t_1/2_) calculated by one phase decay analysis (dote line) from the elimination phase. (**B**) The injected dose of Cy7-psomes after 24 hours i.p. injection in healthy and AIA Wistar rats. Data express the mean ± SEM (n=3 per experimental group).

**Fig. S6.**
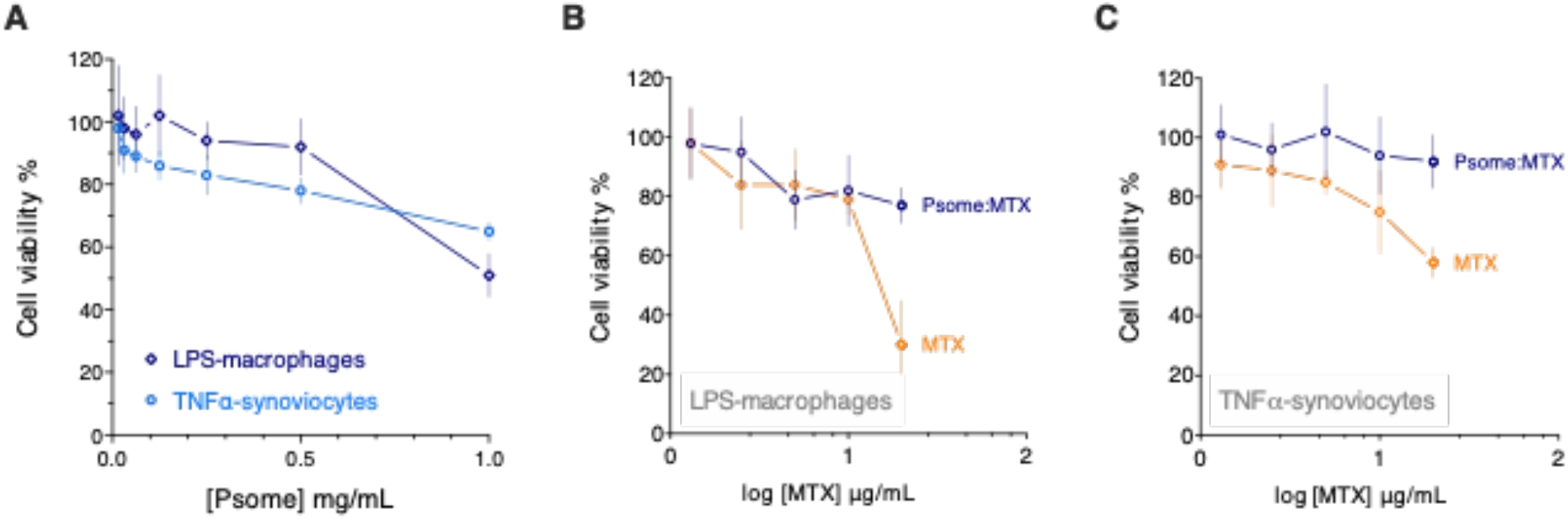
Cell viability of 24 hours treatment with (**A**) psomes, (**B**, **C**) increasing concentrations of either free MTX or MTX-loaded psomes in inflammation-activated macrophages synoviocytes. Data express as mean ± SD (n=3).

**Fig. S7.**
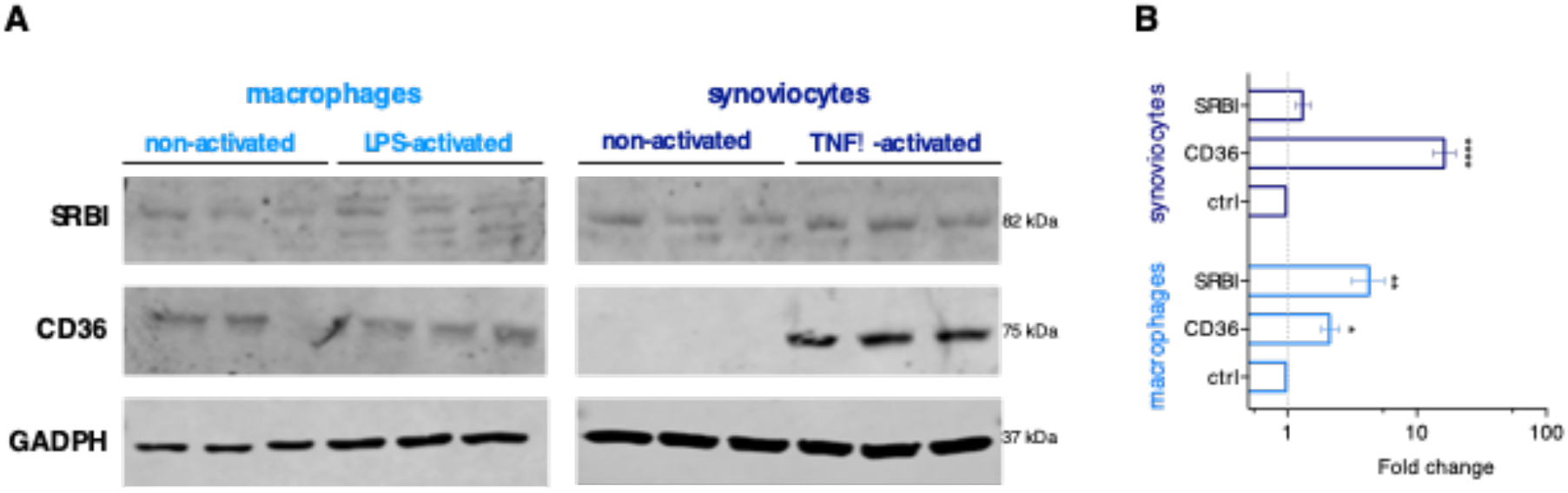
**(A)** SRBI and CD36 cell surface proteins expression levels in non- and activated macrophages and synoviocytes detected by western blot assay (n=3). The SRB1 and CD36 were revealed using specific antibodies and glyceraldehyde 3-phosphate dehydrogenase (GAPDH) was used as the loading control. **(B)** Fold change expression normalised to non-activated control. Data express as mean ± SD (n=3). The differences relative to the non-activated control were statistically significant for *p<0.05, **p<0.01, ***p<0.001 and ****p<0.0001.

**Fig. S8.**
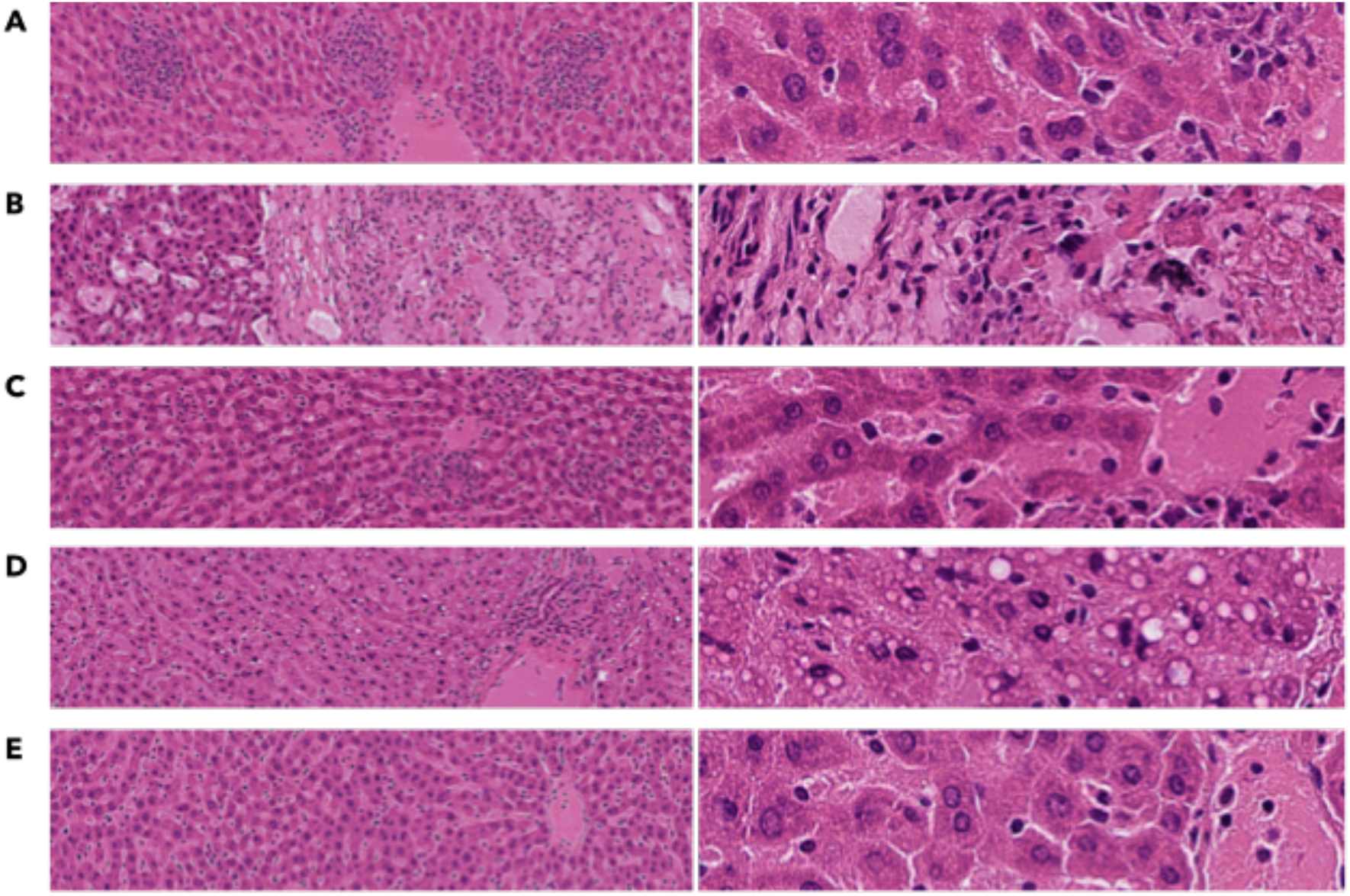
H&E histological representative spleen sections of each experimental animal group: (**A**) healthy; (**B**) AIA; AIA-treated with (**C**) psomes; (**D**) MTX; (**E**) MTX-loaded psomes (left: histology intermediate 5x magnification; right: histology high 20x magnification).

**Fig. S9.**
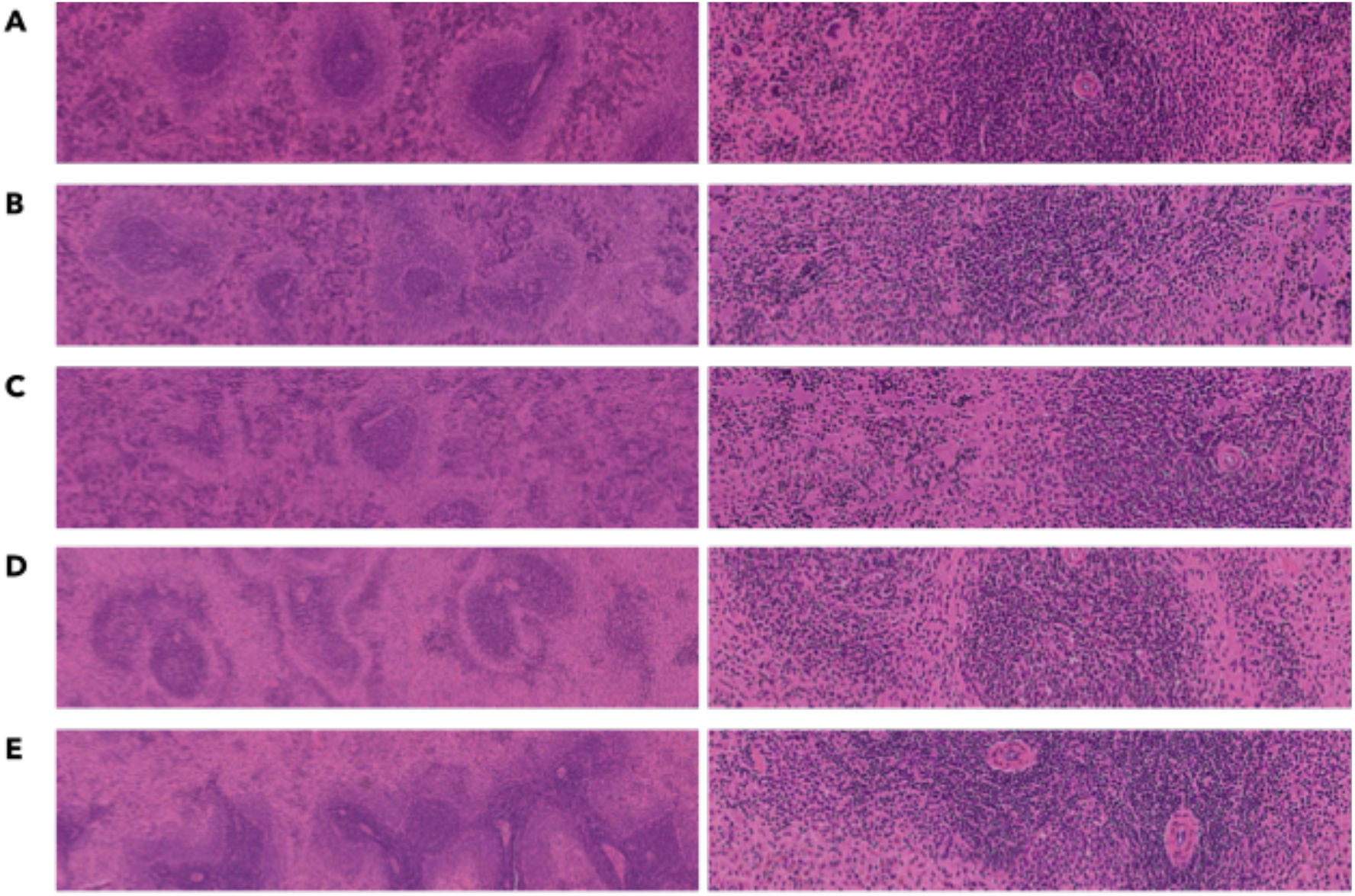
H&E histological representative spleen sections of each experimental animal group: (**A**) healthy; (**B**) AIA; AIA-treated with (**C**) psomes; (**D**) MTX; (**E**) MTX-loaded psomes (left: histology intermediate 5x magnification; right: histology high 20x magnification).

